# Characterisation of the pathogenicity of strains of *Pseudomonas syringae* towards cherry and plum

**DOI:** 10.1101/227223

**Authors:** M.T. Hulin, J.W. Mansfield, P. Brain, X. Xiangming, R.W. Jackson, R.J. Harrison

## Abstract

Bacterial canker is a major disease of cherry and other stone fruits caused by several pathovars of *Pseudomonas syringae*. These are *P.s* pv. *morsprunorum* race 1 *(Psm* R1), *P.s* pv. *morsprunorum* race 2 *(Psm* R2) and *P.s* pv. *syringae* (Pss). *Psm* R1 and R2 were originally designated as races of the same pathovar, however phylogenetic analysis has revealed them to be distantly related. This study characterised the pathogenicity of *P. syringae* on cherry and plum, in the field and the laboratory. The field experiment identified variation in host cultivar susceptibility to the different pathogen clades. The cherry cultivar Merton Glory exhibited a broad resistance to all clades, whilst cultivar Van showed race-specific resistance. *Psm* R1 may be divided into a race structure with some strains pathogenic to both cherry and plum and others only pathogenic to plum. The results of laboratory-based pathogenicity tests were compared to results obtained on whole-trees. Only cut shoot inoculations were found to be sensitive enough to detect cultivar variation in susceptibility. Measuring population growth of bacteria in detached leaves reliably discriminated pathogens from non-pathogens. In addition, symptom appearance discriminated *Psm* races from non-pathogens which triggered a rapid hypersensitive response (HR). The pathogen *Pss* rapidly induced disease lesions and therefore may exhibit a more necrotrophic lifestyle than hemi-biotrophic *Psm* races. This in-depth study of pathogenic interactions, identification of host resistance and optimisation of laboratory assays, will provide a framework for future genetic dissection of virulence and host resistance mechanisms.

## Introduction

*Pseudomonas syringae* is a globally important plant pathogen, and includes strains associated with plants and aquatic environments (Dudnik & Dudler, 2014). Plant pathogenic strains can be divided into pathovars, which are only able to infect particular host species. Strains within pathovars can also be further distinguished into races, which show specificity towards particular host cultivars (Joardar *et al*., 2005). *P. syringae* is referred to as a species complex due to the high level of divergence between individual clades (Berge *et al*., 2014). Currently, nine genomospecies, based on DNA-DNA hybridisation, and thirteen phylogroups, based on Multi-Locus Sequence Typing (MLST), have been described (Gardan *et al*., 1999; Parkinson *et al*., 2011).

Several distantly related pathovars of *P. syringae*, which belong to different phylogroups, are known to cause bacterial canker of *Prunus*. This genus of stone-fruit trees includes economically important species such as cherry, plum, peach and apricot. Focusing on sweet cherry *(Prunus avium)*, members within three distinct phylogroups of *P. syringae* have been characterised as the main causal agents of canker. These are *P. syringae* pv. *syringae* (Pss), *P. syringae* pv. *morsprunorum (Psm)* race 1 (R1) and *P. syringae* pv. *morsprunorum* race 2 (R2) (Bultreys & Kaluzna, 2010). The two *morsprunorum* races are specifically found only on *Prunus* species, whilst *Pss* strain are more variable and able to infect various plant species (Bultreys & Kaluzna, 2010). Although distantly related, *Psm* R1 and R2 were initially distinguished based on virulence towards particular cherry cultivars, so were described as races of pv. *morsprunorum* (Garrett, 1978). Another pathovar (P. *syringae* pv. avii) is pathogenic on wild cherry (Ménard *et al*., 2003). *P. syringae* is able to infect throughout the year and cause necrotic lesions on all aerial plant organs, including fruit, leaves and blossom. The pathovars invade dormant woody tissues through leaf scars and wounds in winter. They occupy the cambial tissue and produce black necrotic cankers in spring. During the growing period, there is a large diverse population of epiphytic bacteria that grow on the surface of the leaves. Bacteria may also enter the leaf and induce necrotic lesions that eventually drop out of the leaf, causing shot-hole symptoms. The asymptomatic leaf population are thought to provide the inoculum for woody tissue infections (Crosse, 1959). Bacterial canker is an annual problem for the global cherry fruit industry and is particularly devastating in young orchards, where it has been reported to cause up to 75% loss of trees (Spotts *et al*., 2010). Chemical control for this disease is currently limited to spraying with copper-based compounds, a treatment that has recently been restricted across Europe (Stone & Baker, 2010). Breeding for resistance is a desirable alternative method of control. Recent studies have identified rootstock selections and scion varieties exhibiting a degree of resistance (Santi *et al*. 2004; Spotts *et al*. 2010; Li *et al*. 2015; Farhadfar *et al*. 2016). Despite this progress, there is still a lack of totally resistant varieties available and the genetic factors underlying canker resistance remain unknown.

An understanding of how the divergent clades of *P. syringae* cause bacterial canker is crucial to breeding efforts. The epidemiology of this disease was determined through field inoculation studies at East Malling in the UK (Crosse, 1966; Crosse & Garrett, 1966; Freigoun & Crosse, 1975; Garrett, 1978). Molecular techniques such as Repetitive Element Sequence-Based (REP) PCR and Multi-Locus Sequence Typing (MLST) and various morphological methods have been used to survey the bacterial populations in orchards (Vicente & Roberts, 2007; Gilbert *et al*., 2008; Kaluzna *et al*., 2010). These studies revealed that the three pathovars co-exist within orchards, with each other and non-pathogenic Pseudomonads. To characterise pathogenicity, several laboratory and field-based assays have been developed (Crosse & Garrett 1966; Vicente & Roberts 2003; Gilbert *et al*. 2009). Improved assays are required for screening for host resistance. Gilbert *et al*. (2009) used various lab-based tests to determine the pathogenicity of strains isolated from stone-fruits in Belgium. They found that no individual laboratory assay could reliably predict pathogenicity under field conditions. Field inoculations are therefore required to fully ascertain pathogenicity and differences in host response.

The breeding of resistant cherry cultivars has been hindered due to the complex nature of this disease (Garrett, 1979). Early work reported variation in cultivar susceptibility towards the different clades of pathogenic *P. syringae*, with two cultivars Napoleon and Roundel exhibiting differential susceptibility towards the two races of *Psm*. Napoleon was found to be resistant to R2 but susceptible to R1, and vice versa for Roundel (Garrett, 1978). It may therefore be challenging to breed resistance to all three of the genetically distinct *P. syringae* clades.

Various studies have established the mechanisms of host immunity in the model *P. syringae* patho-systems of *Solanum lycopersicum* (tomato), *Arabidopsis thaliana* (thale cress) and *Phaseolus vulgaris* (bean) (Preston, 2000; Quirino & Bent, 2003; Arnold *et al*., 2011). *P. syringae* uses a range of virulence factors, including Type III secretion system effector proteins (T3Es) to suppress the plant immune system. Plant immunity can be broadly divided into two stages: PAMP-Triggered Immunity (PTI) and Effector-Triggered Immunity (ETI) (Jones & Dangl, 2006). PTI is a response towards conserved pathogen molecules and allows plants to exhibit non-host resistance to many potential pathogens. ETI occurs when host resistance proteins (R proteins) detect the presence of pathogen effectors and typically leads to a hypersensitive cell death response (HR), which prevents the spread of the pathogen (Senthil-Kumar & Mysore, 2013). ETI is associated with varietal resistance within a host species, whereby particular host cultivars have R genes that trigger the HR towards particular pathogen races. Studies have also found that ETI may play a role in non-host resistance (Gill *et al*., 2015). Initial studies of ETI-associated host resistance were focused on qualitative resistance, whereby a single R gene provides complete resistance against a particular pathogen. Although the above generalisations about PTI and ETI are valid, it is well known that in the field varietal resistance is often quantitative; whereby resistance leads to reduction but not absence of the disease. This type of resistance is often controlled by more than one gene, with different host genotypes exhibiting a range of susceptibility levels to the pathogen (Poland *et al*., 2009), and even towards single T3Es (Iakovidis *et al*., 2016). Quantitative host resistance may encompass a combination of PTI and ETI (Corwin *et al*., 2016). As no fully canker-resistant host genotypes have been identified, the genetic basis of host resistance to bacterial canker is likely to be quantitative. It may involve many different genes that additively contribute to more resistant phenotypes.

To analyse the genetics of pathogenicity and disease resistance to canker, the pathogenicity of a diverse range of *P. syringae* strains was studied. Experiments were conducted, using both lab-based and field inoculations, to characterise the interaction of the strains with both cherry and plum. This robust series of infection experiments provides a reliable pathogenicity framework for future genetic dissection of virulence and disease resistance.

## Materials and Methods

### Bacterial strains

Strains of *Pseudomonas syringae* (listed in Table 1) were grown on King’s B agar at 25 °C. For liquid culture, strains were grown in Lysogeny Broth (LB) shaking at 25 °C, 150 rpm.

**Table 1.**
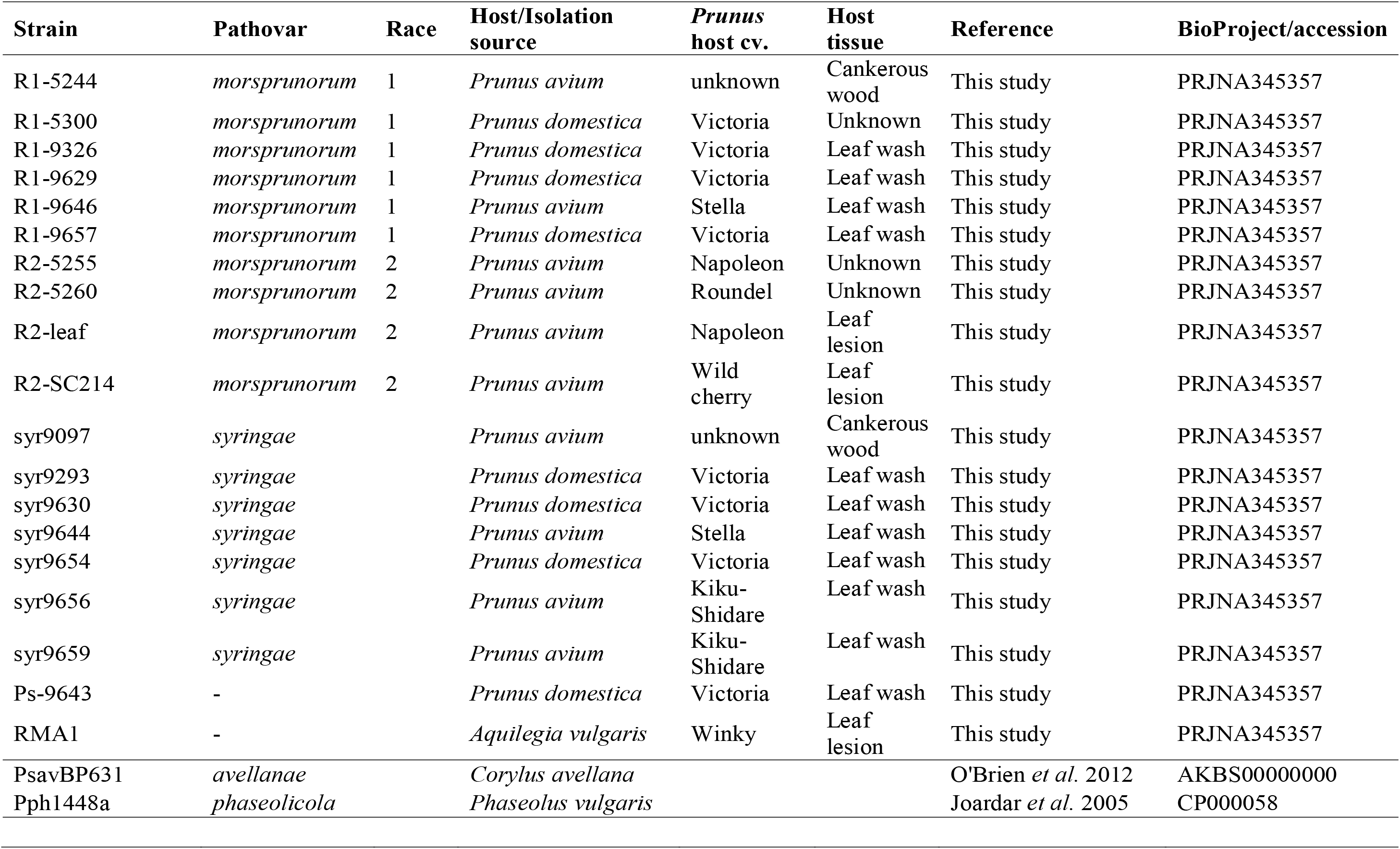
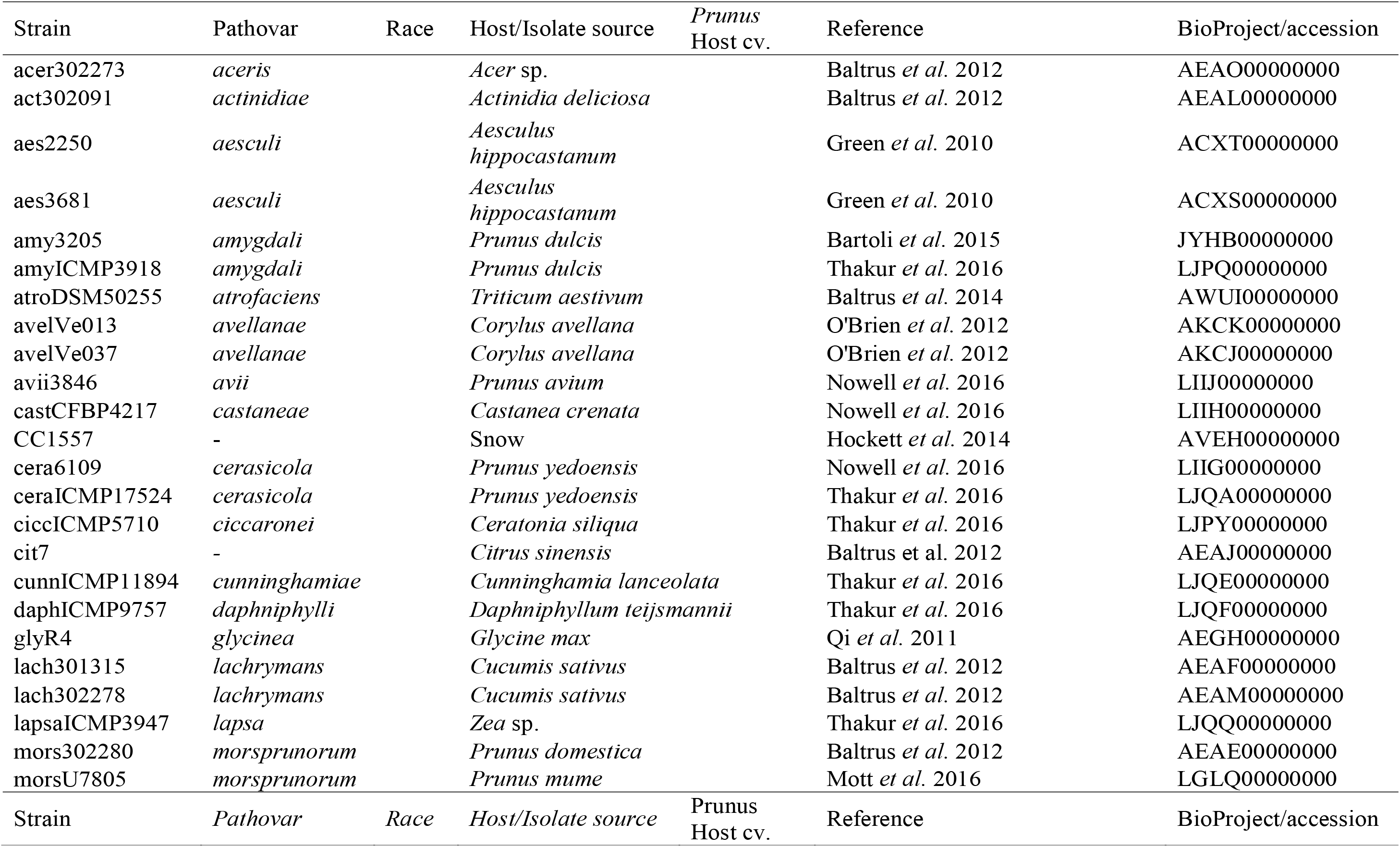
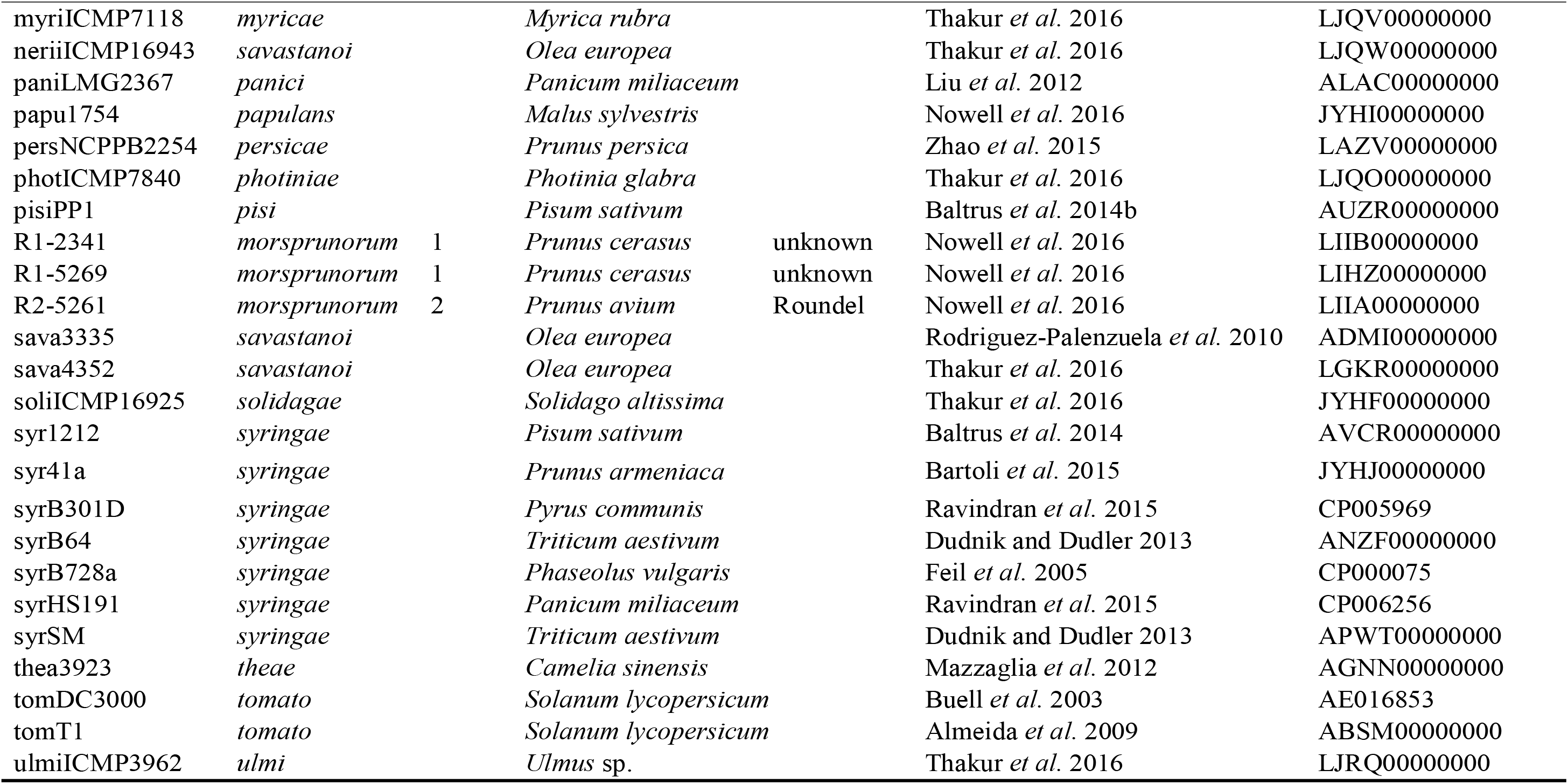
Bacterial strains used in this study with host of isolation and reference/source. Strains sequenced in this study are listed first, followed by the out-group strains *Pph* and *Psav* included in pathogenicity tests and then the rest of the strains used soley for phylogenetic analysis. The Genbank accessions of genomes used for the phylogenetic analysis are included. Full genbank accessions of strains sequenced in this study will be released upon publication.

### Genome sequencing

Nineteen *P. syringae* strains were genome sequenced using the Illumina MiSeq V3. DNA was extracted using the Puregene Yeast/Bact Kit (Qiagen). DNA libraries were prepared by fragmenting the DNA using a sonicating water bath for 30 seconds. DNA was then size-selected by gel electrophoresis to obtain fragments of 400-700 bp using the Zymogen gel extraction kit (Zymo Research). Libraries were created using the NextFlex Rapid-DNA sequencing kit. Barcodes were multiplexed to allow pooling of multiple samples. Libraries were quality checked using the Fragment Analyzer (Advanced Analytical) and Qubit (Life Technologies).

Libraries were sequenced using the Illumina Mi-Seq V3 (Illumina) 300 bp paired-end reads. Raw data for each genome was quality checked and trimmed using fastqc-mcf (https://code.google.com/p/ea-utils/wiki/FastqMcf). The reads were error corrected using Quake prior to assembly (Kelley *et al*., 2010). Each genome was then assembled using SPAdes 3.7.0 (Bankevich *et al*., 2012) and summary statistics generated using Quast (Gurevich *et al*., 2013).

### Phylogenetic analysis

A phylogenetic tree was created for all sequenced strains and other strains which had genome sequences available on NCBI. The nucleotide sequences of seven house-keeping genes *(acnB, fruK, gapA, gltA, gyrB, pgi* and *rpoD*) were extracted from all genomes and individually aligned using Geneious 7.1.9. The alignments were concatenated and trimmed to produce an overall alignment of 9393bp. A Bayesian phylogeny was created of this alignment using the Geneious plug-in of MrBayes (Huelsenbeck & Ronquist, 2001). The GTR gamma model of evolution was used with a burn-in length of 100,000 and sub-sampling frequency of 200.

### Plant material

All *Prunus* material was propagated at NIAB-EMR. For whole-tree inoculations, one-year old grafted trees were used. Cherry cultivars were grafted on the rootstock Gisela 5, whilst plum was grafted on St Julian A. For detached leaf assays, 1-2 week old, fully-expanded leaves were obtained from glasshouse grown trees. Immature green cherry fruits were obtained from mature trees.

### Pathogenicity assays

Bacterial inoculum was prepared from overnight LB cultures. These were spun down (3500 g, 10 minutes) and re-suspended in sterile 10 mM MgCl_2_. A spectrophotometer was used to measure concentration, with an optical density of 0.2 (OD_600_) being ~2x10^8^ CFU/ml (Debener *et al*., 1991).

### Whole-tree inoculations

Whole-trees were inoculated though either wounds or leaf scars (Crosse & Garrett, 1966). Field inoculations were performed in October 2015 and glasshouse wound inoculations in February 2015. Bacterial suspensions of 2x10^7^ CFU/ml were used for inoculations.

To inoculate through wounds, a sterile scalpel was used to cut a shallow wound into the trunk of the tree and 200μl of inoculum was pipetted into the wound. To inoculate leaf scars, the leaf was removed and 10μl of bacterial suspension was pipetted on the exposed scar. The inoculation sites were covered with parafilm and duct tape. Multiple inoculations were performed on the same tree, with at least 4 buds between inoculations. For the field experiment the trees were left for 6 months before assessment in May 2016, whereas the glasshouse experiment was assessed after 2 months.

To assess disease the bark was stripped back and length of necrosis was measured using a caliper (field experiment only). A disease score was determined as 1: no symptoms 2: limited browning, 3: necrosis and gumming and 4: necrosis, gumming and spreading from site of inoculation.

### Cut shoot inoculations

A cut shoot assay was performed as in previous studies (Krzesinska *et al*., 1992; Santi *et al*., 2004). Eight strains were inoculated onto four cherry and two plum cultivars. Bacterial inoculum was prepared at a concentration of 2x10^7^ CFU/ml. Dormant one-year shoots (5mm diameter) were cut into 10 cm sections. They were sterilised with 0.5% hypochlorite for five minutes, rinsed in tap water and left overnight to air-dry. Next, 5 mm from the shoot tip was cut and the shoot was dip inoculated for five minutes. The wound was covered with parafilm and the shoot bases were freshly cut (5 mm) and placed in transparent-boxes immersed in 20 mm deep distilled water. The shoots were incubated at 15 °C with 16-hour light, 8-hour dark cycle for one week. Next, shoots were transferred to −2 °C for one week to simulate frost damage. Finally, the basal 10 mm of each shoot was removed and they were placed in water-soaked Oasis Foam (Oasis Floral). These were incubated for a further 4 weeks at 15 °C. The trays were covered with cling-film to maintain a high humidity.

The shoots were assessed by peeling back the bark from the top 30 mm of the shoot. Digital images were captured and analysed with software (Li *et al*., 2015) to determine the percentage area of necrosis.

### Cherry fruit inoculations

To inoculate immature cherry fruits a stab-inoculation method was used (Moragrega & Llorente, 2003). Fruits were sterilised in 0.5% hypochlorite for five minutes and rinsed in distilled water. Bacteria were then scraped from 5-day old plates using a 24g needle and stabbed into the plant material. Fruits were placed in transparent boxes lined with moist tissue paper to maintain a high humidity. The fruits were kept at 22 °C (16hr light, 8hr dark) and visually assessed over time. Two independent assays on the different cultivars were performed.

### Leaf inoculations and microscopy

Inoculum concentration varied from 2x10^6^ CFU/ml to 2x10^8^ CFU/ml. Freshly picked, 1-2 week old leaves were used for leaf inoculations. The leaves were infiltrated with bacterial suspension from the abaxial surface using a blunt-ended syringe. Leaves were then placed in plastic trays, which contained a 10 mm layer of water agar (10g/L), covered in damp paper towel. The tray was sealed inside a transparent bag and incubated at 22 °C (16hr light, 8hr dark). The leaves were left for a maximum of 10 days before assessment. At least three leaves were inoculated for each isolate, with the three replicate leaves coming from different plants.

Bacterial population growth within the leaves was measured over time. Day 0 populations were always calculated to check that the inoculum concentrations were similar between treatments. Leaf discs were excised using a sterile cork borer (0.5cm). Discs were then homogenised in 10 mM MgCl_2_. A dilution series was plated out to determine bacterial concentration (CFU/ml). Each concentration was plated out three times (pseudoreplicates). Overall, for each bacterial strain studied there were three replicate leaf inoculations and three pseudoreplicates to measure the concentration of each. Two independent experiments were performed for the leaf assays of a subset of strains inoculated on cherry and plum and the inoculations on different cherry cultivars.

Electron microscopy was performed by Dr Ian Brown (University of Kent) on infected cherry leaves. Detached leaves were infiltrated with bacteria at 2x10^6^ CFU/ml and incubated for one week at 22 °C.

Inoculated leaves were cut into 2 mm squares using a razor blade in a drop of cold fixative (2.5% glutaraldehyde in 100mM sodium cacodylate buffer pH 7.2 [CAB]) on dental wax and processed as previously described (Soylu *et al*., 2005). Sections were viewed in a Jeol 1230 TEM with an accelerating voltage of 80kV and images recorded with a Gatan Multiscan 791 digital camera.

### Experimental design and statistical analysis

To randomise the glasshouse whole-tree canker assay, an incomplete block-design was used. This allowed assessment of 22 different strains on 22 trees, with five replicates of each strain. In the field experiment, the virulence of eight strains was assessed on four cherry and two plum cultivars, using two different inoculation methods. To reduce the number of trees required, the eight different strains were divided across two trees, with each tree also having one negative control. This meant that two adjacently planted trees comprised one experimental unit of all strains and controls inoculated on the same cultivar using one inoculation method. A balanced incomplete design was used to randomise strain positions onto the two trees. A balanced complete design was then used to randomise the different cultivars and inoculation methods within 10 blocks in the field. Each block contained 24 trees (16 cherry and eight plum), and the total experiment involved 240 trees.

R software (R Core Team, 2012) was used for all statistical analyses as described in detail in supplementary methods. All ANOVA tables are also presented in the supplementary data.

## Results

### Phylogenetics

To determine the diversity of strains isolated from cherry and plum, the genomes of 18 *P. syringae* strains were sequenced. The strains included bacteria representative of all three previously designated clades, *Psm* R1, *Psm* R2 and *Pss. Pss* and *Psm* R1 included strains isolated from both cherry and plum, whilst the *Psm* R2 strains all originated from cherry. A previously undescribed strain that did not belong to these clades *(Ps* 9643), which had been isolated from a plum leaf wash was also included. Finally, an additional strain (RMA1) isolated from the perennial species *Aquilegia vulgaris*, that preliminary analysis had shown to be closely related to *Psm* R2 was sequenced. The DNA sequences of seven MLST genes were then extracted from the genomes. Homologous sequences from 59 genome assemblies of additional strains within the *P. syringae* complex were then downloaded from NCBI. These included strains that were also isolated from *Prunus* and other plant species for comparison. A Bayesian phylogenetic tree was then generated based on a concatenated alignment of the seven genes.

The *P. syringae* phylogeny is presented in Figure 1, with strains isolated from *Prunus* highlighted. *Psm* R1, *Psm* R2 and *Pss* were found within phylogroups 3, 1 and 2 respectively. The two *Psm* races fell into discrete monophyletic clades, with individual strains being very closely related. By contrast, *Prunus Pss* isolates exhibited greater diversity. Strains isolated from cherry and plum did not form distinct host-specific clusters in any of the pathogenic clades, indicating that they are closely related and may cross-infect the two *Prunus* species. *Ps* 9643 was closely related to the *Prunus persicae* pathogen *(P.s* pv. *persicae)*, whilst RMA1 was an out-group to the clade containing *Psm* R2 and the pathovars *P.s*. pv. *actinidiae, P.s* pv. *avellanae* and *P.s*. pv. *theae* (which infect kiwifruit, hazelnut and tea respectively).

**Figure 1.**
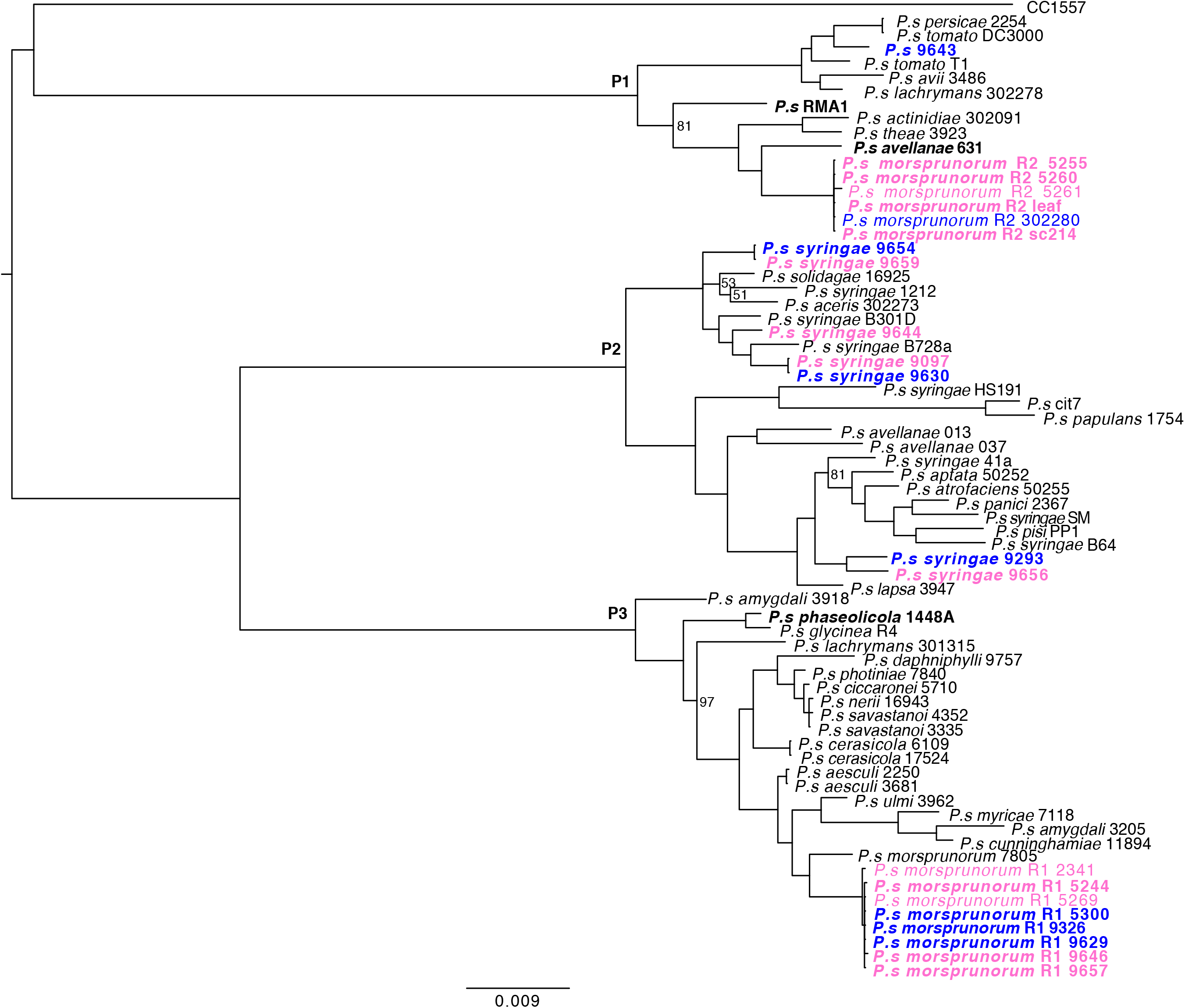
Bayesian phylogenetic tree of *P. syringae*. The phylogeny was constructed using a concatenated alignment of seven genes (*acnB, fruK, gapA, gltA, gyrB, pgi* and *rpoD*). A subset of strains from the three major phylogroups were selected for analysis, with the canker-causing clades *Psm* R1, *Psm* R2 and *Pss* highlighted. Phylogroups are labelled P1-3. Strains isolated from cherry are in pink, whilst those from plum are in blue. Strains in bold were pathogenicity tested in this study. Scale bar shows substitutions per site. Bootstrap support values <99% are presented.

### Characterising pathogenicity of a range of *P. syringae* strains on cherry and plum trees

#### Whole-tree glasshouse experiment

To determine the fundamental ability of each strain to cause bacterial canker on cherry, a whole-tree wound inoculation experiment was performed. All strains isolated from cherry and plum, as well as related pathogens of other plants: *P.s* pv. *phaseolicola* 1448A *(Pph), P.s* pv. *avellanae* BPIC631 *(Psav)* and RMA1, were included. Comparisons between strains were made based on the level of necrosis produced in the cambial layer underneath the bark at the site of inoculation after two months of incubation. Strains exhibited a wide range of virulence profiles on cherry (Figure 2). Both the non-pathogens and negative control gave very limited browning and callusing associated with a wound response. Pathogenicity was indicated by black, necrotic lesions that sometimes spread from the inoculation site and were associated with gumming. There was clear variation between members of the different *Prunus-infecting* clades. Strains of *Psm* R1 and R2 showed variation in virulence, but rarely spread from the inoculation site. Meanwhile, most strains of *Pss* were able to spread. Within *Psm* R1, only two cherry strains (R1-5244 and R1-9646) caused gumming and necrosis, whilst R1-9657 showed reduced virulence, not significantly different to the plum R1 strains. Symptoms caused by strains of *Psm* R1 isolated from plum were not significantly different from those associated with the non-pathogens. Most strains of *Psm* R2 were pathogenic, however R2-5260 showed reduced virulence. Apart from one strain, *Pss* was highly pathogenic, with symptoms typically spreading from the site of inoculation. The strain *Ps* 9643 isolated from a plum leaf, but found not to be closely related to the other canker pathogens (Figure 1), behaved as a non-pathogen of cherry.

**Figure 2.**
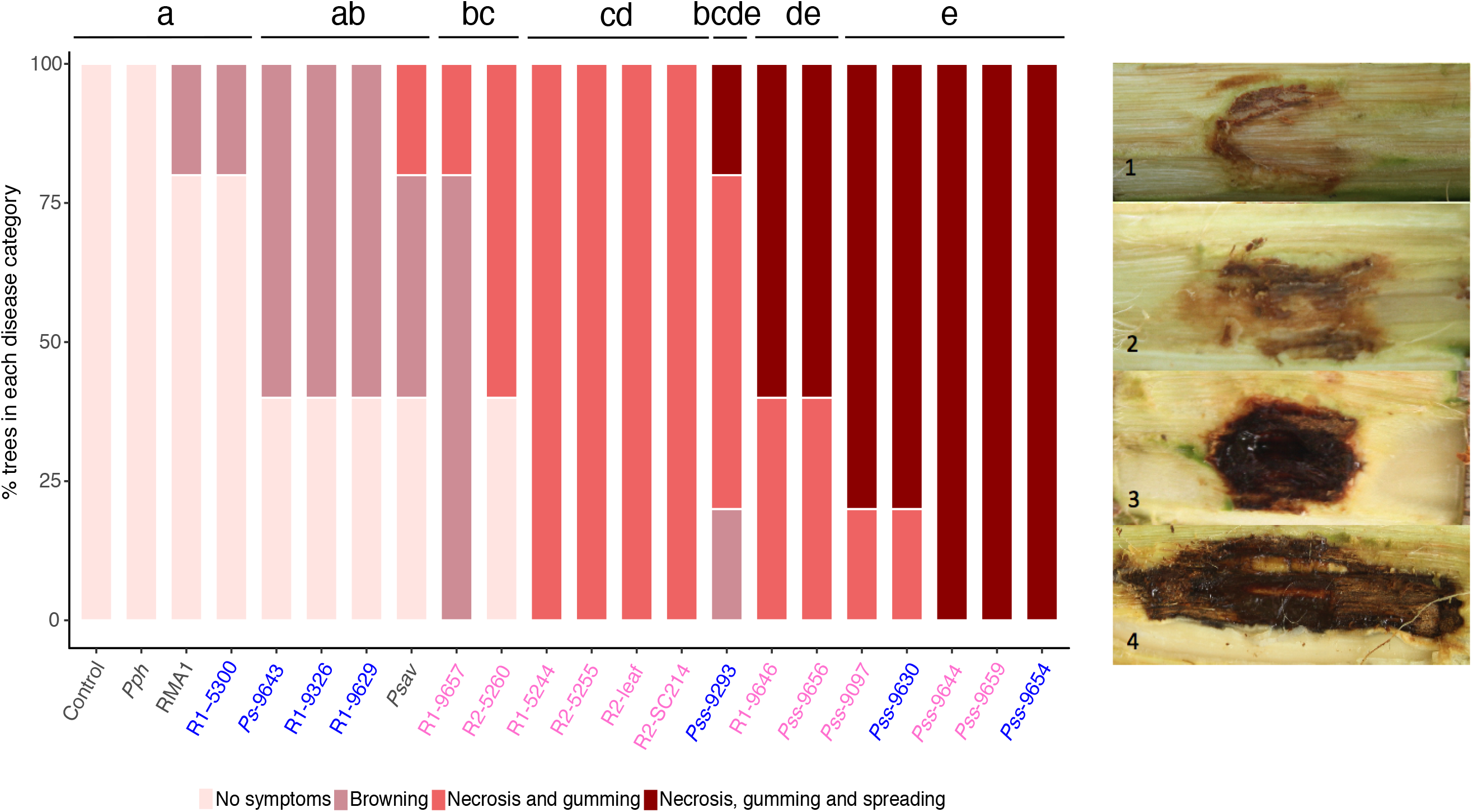
Percentage of trees in each disease score category after wound inoculation of *P. avium* cv. Van with different isolates of *P. syringae: Psm* R1, *Psm* R2, *Pss*, selected outgroup non-host strains and a no bacteria control. Data presented are the percentage of replicates (n=5) for each strain in each disease category. Disease symptoms were scored on a ordinal scale as illustrated: 1, no symptoms; 2, limited browning; 3, necrosis and gumming; 4, necrosis, gumming and spread from site of inoculation. Strains are ordered based on increasing disease score. Strains isolated from cherry are labelled in pink, whilst those from plum are in blue. Statistical Tukey-HSD (p=0.05, confidence level: 0.95) groupings of bacterial strains determined by a Proportional Odds Model (POM) analysis are presented above the bar.

#### Whole-tree field experiment

A set of strains with contrasting pathogenicity and host of isolation was chosen for pathogenicity screening under field conditions, using leaf scar and wound inoculations on cherry and plum cultivars. The strains included cherry pathogens (R1-5244, R2-leaf, *Pss* 9097 and *Pss* 9293) and non-pathogens (R1-5300, *Ps* 9643, *Pph* and RMA1). Concerning cherry, the cultivar Merton Glory is reported to be tolerant to canker (APS, 1966), Napoleon and Roundel show race-specific differences, with Napoleon being susceptible to *Psm* R1 and tolerant to R2 (and vice versa in Roundel) (Garrett, 1978). The cultivar Van is reported to be universally susceptible (Long & Olsen, 2013). For plum, Victoria is reported as susceptible and Marjorie’s Seedling is more resistant (RHS, n.d.).

In cherry, data for both disease score (on an ordered categorical scale) and symptom length (mm) are presented in Figure 3. With both inoculation methods, the pathogens (R1-5244, R2-leaf, *Pss-* 9097 and Pss-9293) caused necrosis and gumming (score ≥3), and in some cases lesions spread extensively beyond the inoculation site. In contrast to the glasshouse wound inoculations, all three pathogenic clades *(Psm* R1, *Psm* R2 and *Pss)* were able to spread from site of inoculation (previously only *Pss* appeared to spread). The non-pathogen inoculations generally induced limited browning (scores 1-2), with disease score profiles similar to the control. In the field, contamination by wild Pseudomonads may have occurred, and this explained why some control inoculations generated disease symptoms (6% of controls scored ≥3). For disease score, both inoculation methods were analysed together. The percentage of inoculations exhibiting disease symptoms (score ≥ 3) was greater in the wound inoculations than scar. Whilst, comparing cultivars, higher scores were more frequently observed in Napoleon than the other three cultivars.

**Figure 3.**
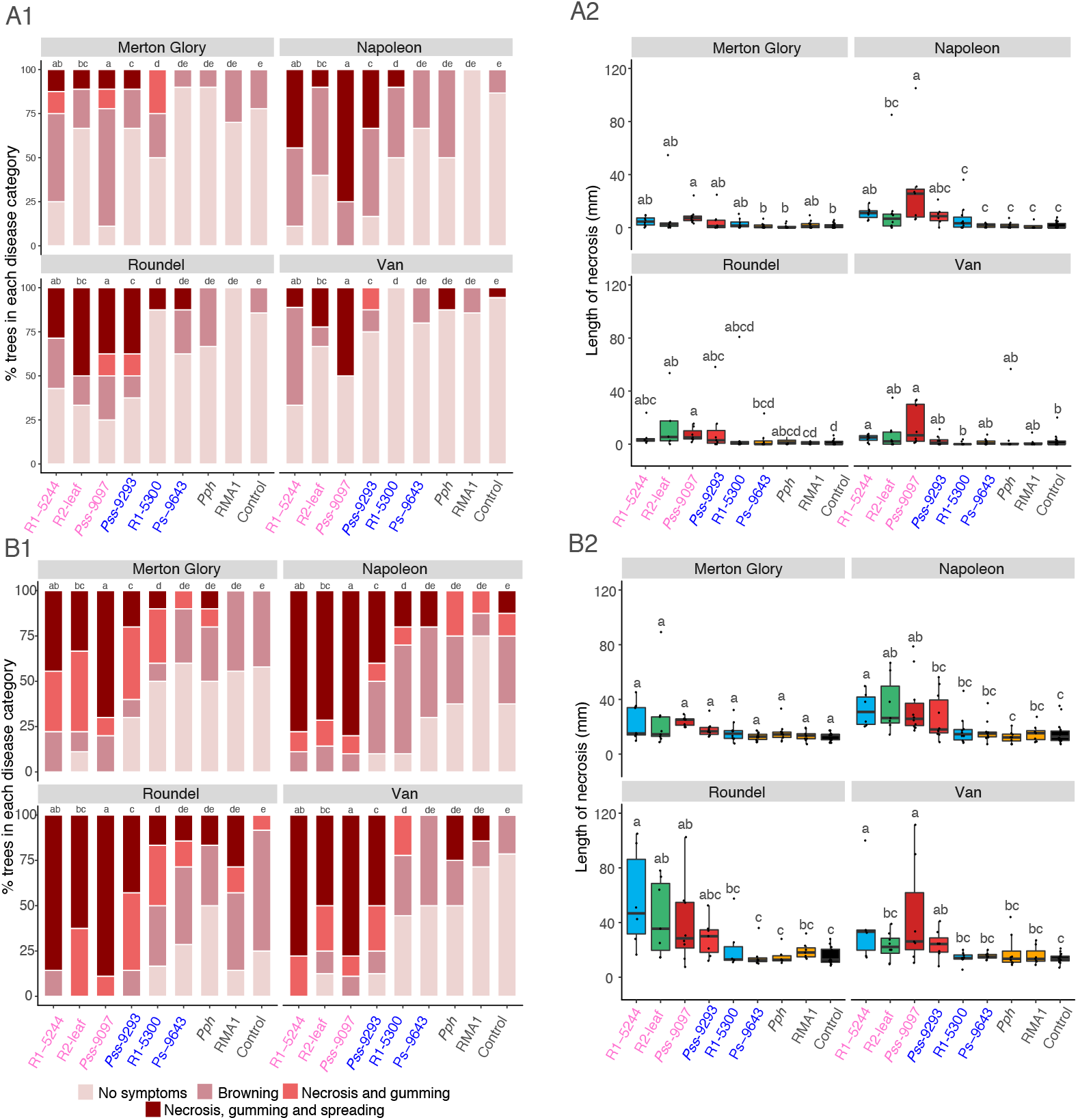
Field inoculations of different cherry cultivars with selected strains of *P. syringae*. Data presented are the disease score and length of disease symptoms based on symptom observations six months after inoculation. A: Leaf scar inoculation. B: Wound inoculations. 1: Percentage of trees in each disease score category (colour-coded from light pink to red by score: no symptoms, browning, necrosis and gumming and necrosis, gumming and spreading from site of inoculation). 2: Boxplot of length of symptoms associated with each strain on the four cultivars. Boxplots are colour-coded for each strain based on clade *Psm* R1 (blue), *Psm* R2 (green), *Pss* (red), outgroup avirulent strains (orange) and no bacterial control (black). All data points (n=10) are presented. Strains on all plots are colour-coded based on host of isolation (cherry in pink, plum in blue and other hosts in black). For disease score POM analysis indicated that there was a significant difference between inoculation method (p<0.01, df=1), between *P. syringae* strains (p<0.01, df=8) and between cultivars (p<0.01, df=3). For symptom length, REML analyses indicated there were significant differences between strains and cultivars for both the leaf scar and wound experiments (p<0.01, df=8 and p<0.01, df=3 respectively). Tukey-HSD (p=0.05, confidence level: 0.95) groups are presented above each strain for each cultivar.

Data for lesion length are also presented in Figure 3 (A2/B2). Due to differences in variance the two inoculation methods were analysed separately. REML analyses indicated there were significant differences between bacterial strains and host cultivars for both inoculation experiments. In both the length and score analyses there were no significant interactions between treatments, as pathogen and non-pathogen responses were consistent across the cultivars, with the two inoculation methods. There did appear to be variation in *Psm* R2 virulence between the cultivars, particularly after wound inoculation (Figure 3-B2), with reduced virulence compared to *Psm* R1 on Van, but a high level of virulence on Roundel. The plum strain *Psm* R1 5300 was not significantly different from the nonpathogens, indicating that it lacks pathogenicity for cherry. The two *Pss* strains varied considerably in virulence, with the cherry isolate *Pss* 9097 being associated with higher disease scores than the plum isolate *Pss* 9293. This is consistent with the results of the glasshouse inoculation, where *Pss* 9293 showed a reduced ability to cause canker. The cultivar Merton Glory appeared to be more tolerant to canker, with the lowest overall mean symptom length. Pathogenic strains were able to cause disease symptoms on this cultivar but the length of these symptoms were not significantly greater than non-pathogens. This indicated that the pathogens could cause disease, but not spread effectively from site of inoculation.

In plum (Figure 4), symptoms produced were similar to those on cherry, with necrosis and gumming being indicative of disease. For the disease score, only strains with confirmed pathogenicity were able to spread (score = 4), however in comparison to the cherry inoculations, the R1-5300 plum isolate was pathogenic. As in cherry, infections through scars produced reduced disease scores compared to wound inoculations. The lesion length analysis showed that the only strains that were ever significantly different from the control were *Pss* 9097 and *Psm* R1 5300. Although analysis revealed there was no significant difference between the two cultivars, the plum cultivar Marjorie’s Seedling did not appear to be susceptible to leaf scar infection, as no strain caused a necrosis length significantly different from the control. The plum cultivar Victoria was slightly more susceptible, with all strains of *Pss* and *Psm* R1 causing some necrosis.

**Figure 4.**
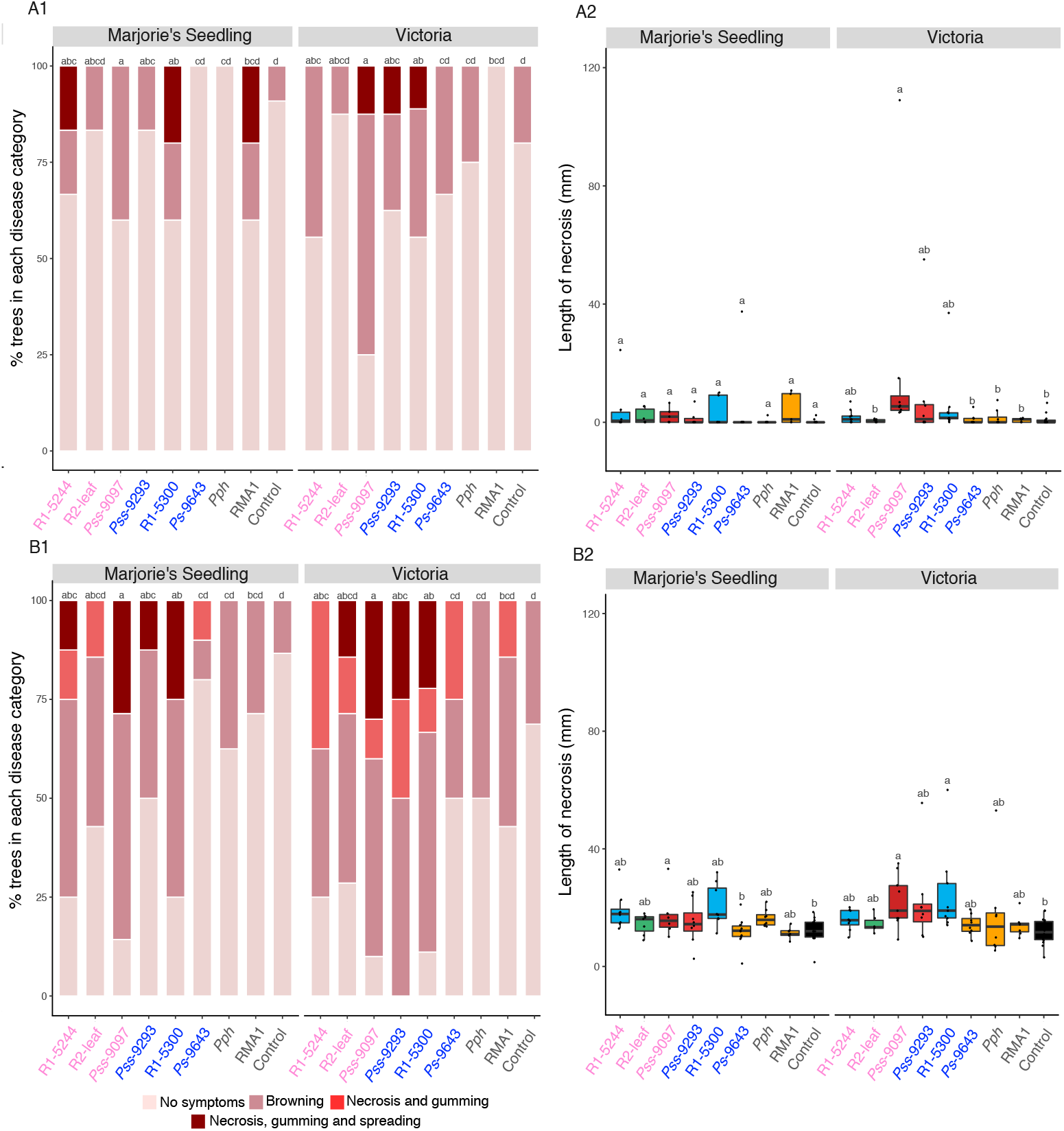
Field inoculations of different plum cultivars with selected strains of *P. syringae*. Data presented are the disease score and length of disease symptoms based on symptom observations six months after inoculation. A: Leaf scar inoculation. B: Wound inoculations. 1: Percentage of trees in each disease score category (colour-coded from light pink to red by score: No symptoms, browning, necrosis and gumming and necrosis, gumming and spreading from site of inoculation). 2: Boxplot of length of symptoms associated with each strain on the four cultivars. Boxplots are colour-coded for each strain based on clade *Psm* R1 (blue), *Psm* R2 (green), *Pss* (red), outgroup avirulent strains (orange) and no bacterial control (black). All data points (n=10) are presented. Strains on all plots are colour-coded based on host of isolation (cherry in pink, plum in blue and other hosts in black). For disease score POM analysis indicated there were significant differences between inoculation method (p<0.01, df=1), strains (p<0.01, df=8) and cultivars (p<0.01, df=1). For symptom length, REML analyses indicated there were significant differences between strains in both inoculation experiments (p<0.01, df=8) but not between host cultivars (p=0.20, df=1 for leaf scar, p=0.35, df=1 for wound). Tukey-HSD (p=0.05, confidence level: 0.95) groups are presented above each strain for each cultivar.

### Laboratory-based pathogenicity assays

The whole-tree inoculations allowed the the virulence of different *P. syringae* strains to be determined and identified non-pathogenic isolates. To rapidly screen for differences in virulence these methods are slow and involve the destruction of whole trees. To undertake large-scale resistance screens of *Prunus* mapping populations or to perform molecular studies of pathogenicity these methods are intractable. Therefore, several laboratory-based assays were assessed for their ability to reflect infection of whole trees.

### Cut shoot inoculations

Several studies have documented the use of detached shoots for screening for bacterial canker resistance (Krzesinska *et al*., 1992; Santi *et al*., 2004; Li *et al*., 2015). Using strains included in the field assay, cherry and plum were screened with the cut shoot method. This involved using one-year old dormant shoots and inoculating a cut end by dipping in bacterial suspension. The extent to which necrosis spread down the shoot cambial tissue from this point could then be used to measure quantatitive differences in bacterial virulence/host resistance.

Figure 5 presents the results on both cherry and plum shoots. Strains exhibited host specificity towards the two *Prunus* species and towards particular cultivars. Focusing on cherry, pathogenic strains within *Psm* R1-5244, *Psm* R2-5255 and *Pss* 9097 were able to cause necrosis on >5% of the shoot area. The two *Psm* races varied in virulence on the different cultivars. As in the field experiment, *Psm* R2 was more virulent on Roundel, but less virulent on Van compared to *Psm* R1. The cut shoot test also confirmed that Merton Glory showed some tolerance compared to the other cultivars. On plum, the level of necrosis on Victoria was greater than that on Marjorie’s Seedling. As observed in the field experiment, the plum strain of *Psm* R1 (R1-5300), was able to cause necrosis where it had failed on cherry. On cv. Victoria the *Aquilegia* pathogen RMA1 caused necrosis similar to *Pss* 9097.

**Figure 5.**
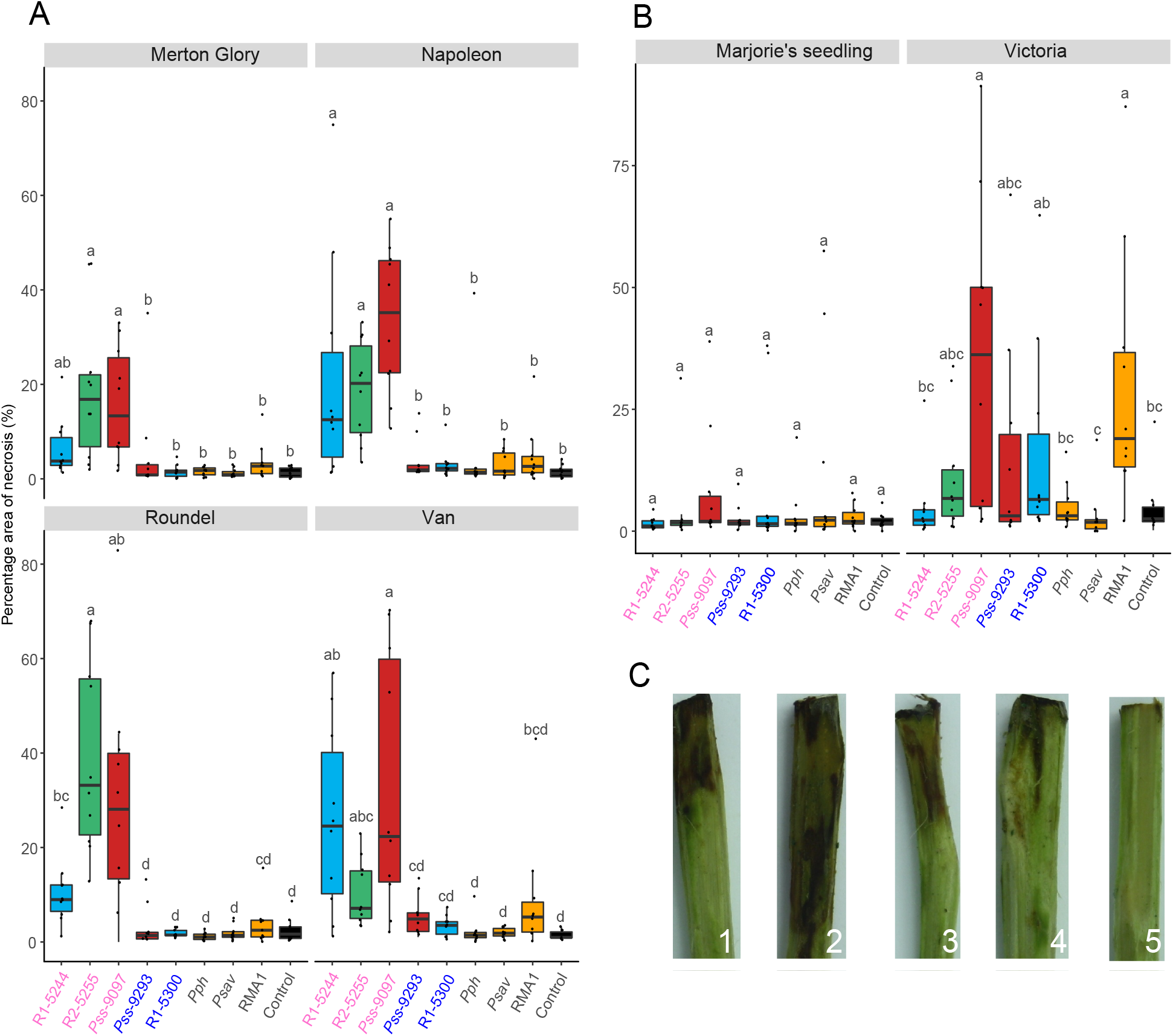
Lesion development on cut shoots of cherry and plum cultivars following inoculation with *P. syringae*. A: Boxplot of percentage area of necrosis in the top 30mm associated with different *P. syringae* strains on four cherry cultivars. All data points for each treatment (n=10) are presented. The bar chart is colour-coded based on clade, *Psm* R1: blue, *Psm* R2: green, Pss: red, non-pathogens: orange and control: black. B: The same parameters for two different plum cultivars. C: Representative images of the symptoms on shoots inoculated with *Pss* 9097 on cv. Napoleon (1-4) or the no bacteria control (5). Strain numbers on all plots are colour-coded based on host of isolation (cherry in pink, plum in blue and other hosts in black). An ANOVA revealed there were significant differences between bacterial strains (p<0.001, df=8), no significant difference between the susceptibility of the two *Prunus* species (p=0.57, df=1) and there was a significant interaction between *Prunus* species and *P. syringae* strain (p<0.01, df=8) as well as interactions between strain and individual cultivars (p<0.01, df=36). Tukey-HSD (p=0.05, confidence level: 0.95) significance groups for the different strains for each separate cultivar are presented above each boxplot.

### Inoculation of detached immature cherry fruits

The suitability of immature cherry fruits was assessed for screening for bacterial canker resistance. Following stab inoculation, symptoms developed within a few days. Examples of the different clades that infect *Prunus* produced remarkably different symptoms on this tissue. Strains of *Pss* produced large necrotic lesions on cherry fruits within 2 days, and these expanded over time. By contrast, both *Psm* races produced water-soaked lesions within 2 days, and these did not increase in size. Most of the non-pathogens caused limited browning. Qualitative symptom assessment therefore allowed differentiation between pathogens and non-pathogens (Figures S1-S4). Measurements of lesion diameter caused by all *P. syringae* strains (Figure 6), confirmed significant differences between strains. However, diameters of the *Psm*-induced water-soaked lesions were not greater than non-pathogens symptoms. The strain *Ps* 9643, although non-pathogenic on trees, caused a similar level of water-soaking to the pathogenic *Psm* races (Figure S4), indicating a failure of fruit to mount a resistant reaction towards this strain.

**Figure 6.**
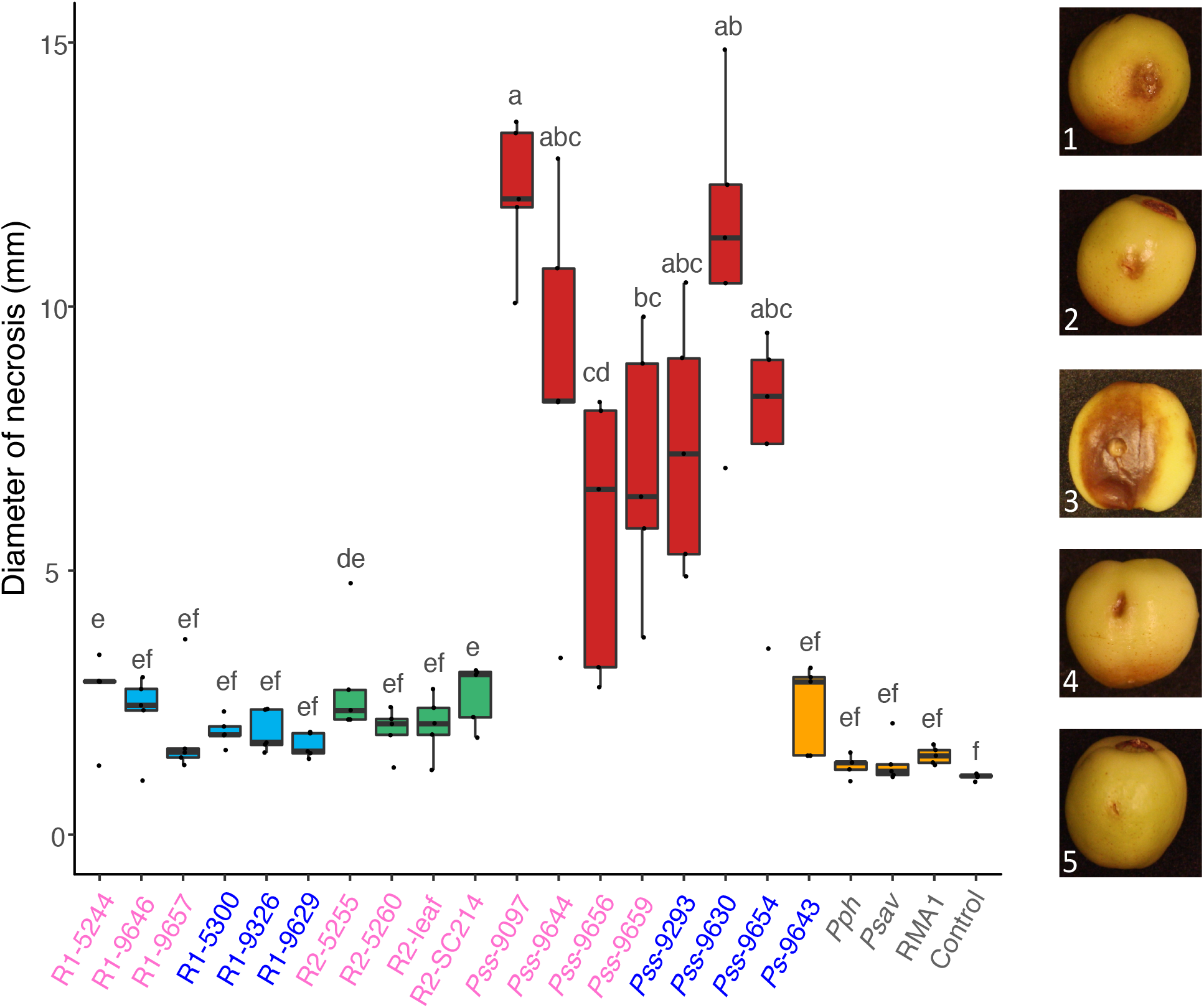
Boxplot to show diameter of necrosis caused by different *P. syringae* strains on immature cherry fruits. Strains isolated from cherry and plum are highlighted in pink and blue. The bar chart is colour-coded, *Psm* R1: blue, *Psm* R2: green, *Pss:* red, non-pathogens: orange and control: black. All data points for each treatment (n=5) are shown. Representative images are presented. 1: *Psm* R1, 2: *Psm* R2, 3: *Pss*, 4: non-pathogens, 5: control. An ANOVA revealed significant differences between strains (p<0.01, df=21). Tukey-HSD (p=0.05, confidence level: 0.95) significance groups are presented above each bar.

The three pathogenic clades were then used to screen different cherry cultivars. The results for lesion diameter are presented in Figure S5. Lesions caused by *Pss* were smaller on Merton Glory and Napoleon than on Van. However, no differences in lesion size or appearance between the two *Psm* races on different host cultivars were found, in contrast to experiments on woody tissues.

### Inoculation of detached leaves

A pilot experiment determined the best method of leaf inoculation was by blunt syringe-infiltration (Figure S6). Bacterial multiplication was initially recorded following inoculation with a low concentration of bacteria (2x10^6^ per ml). On cherry, the pathogens (R1-5244, R2-leaf and *Pss-* 9097) exceeded levels of 10^6^ CFU/ml within four days (Figure 7) and caused black necrosis at the site of infection. The non-pathogens, including the plum isolate R1-5300 failed to reach 10^6^ CFU/ml even after 10 days *in planta* and did not produce symptoms. These results support those found on whole-trees, with only those strains capable of causing bacterial canker being able to reach high levels within leaves. On plum, the pathogens also exceeded 10^6^ CFU/ml after 4 days. However, some of the strains that were non-pathogenic on cherry were able to grow to similar levels as the pathogens. The *Psm* R1 plum isolate R1-5300 and RMA1 isolated from *Aquilegia vulgaris* were found to be capable of multiplication. In the case of R1-5300 this result supports results from inoculation of woody tissues indicating that it is a pathogen of plum but not cherry. The ability of RMA1 to multiply within plum leaves did not support the field experiment where it caused similar symptoms to the negative control.

**Figure 7.**
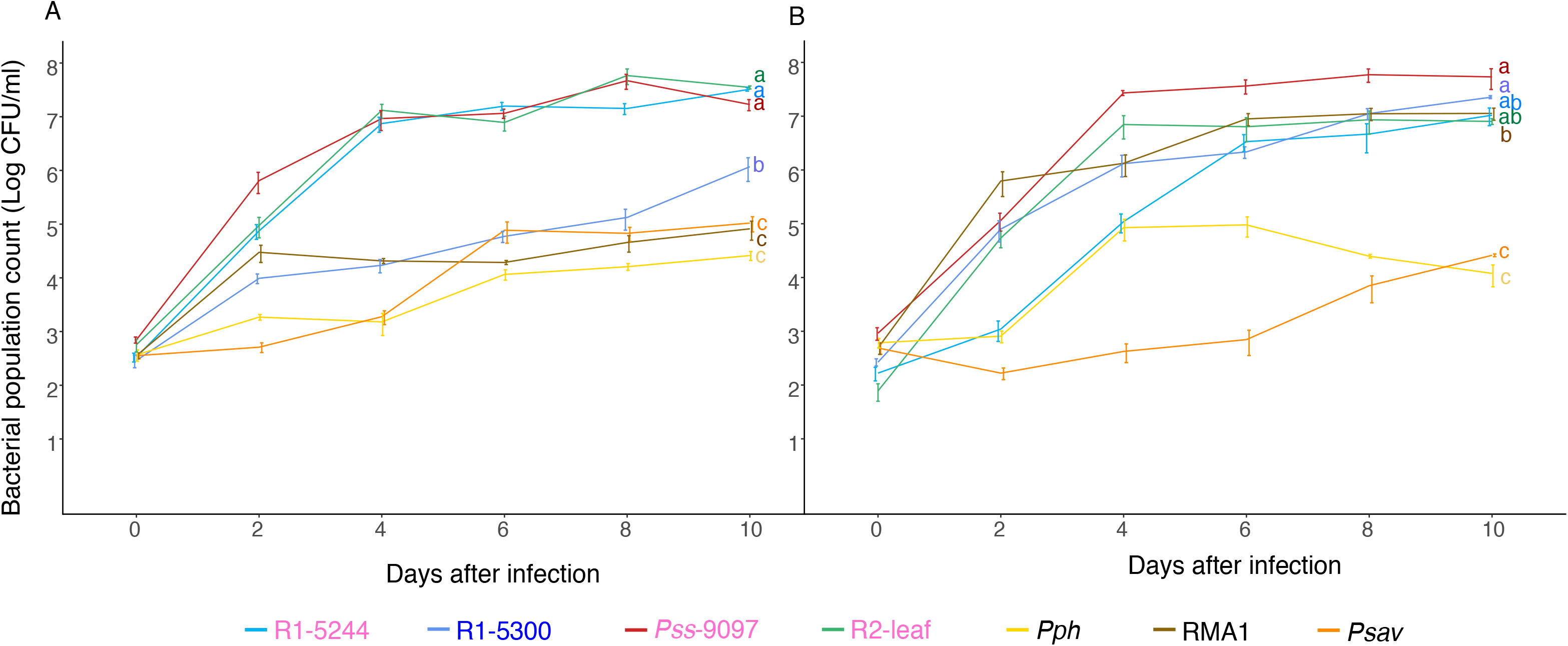
Population counts of different strains over time on cherry cv. Van (A) and plum cv. Victoria (B) leaves. The strains isolated from cherry and plum are highlighted in pink and blue. Line colours for each strain are presented in the key. Population counts are Log CFU/ml. Data presented are the mean values (n=9), with error bars showing standard error above and below the mean. An ANOVA revealed significant differences between strains (p<0.01, df=8).Tukey-HSD (p=0.05, confidence level: 0.95) significance groups for the different strains (based on day 10 populations) are presented.

Next, the population growth of all strains used in this study was tested. An end-point bacterial population count was taken after 10 days. Statistical analysis grouped the pathogens and non-pathogens into separate groups, validating population measurements as a method to differentiate pathogenic and non-pathogen strains (Figure S7).

The induction of the HR was also tested. To determine the best concentration to detect a HR, symptom development was scored at different concentrations (Figure S8). Scores were 0: no lesion, 1: limited browning, 2: browning <50% of inoculated area, 3: browning >50% of inoculated area, 4: complete browning, 5: browning and spread from inoculation site. Area Under the Disease Progression Curve (AUDPC) values were calculated to make comparisons based on timing of symptoms. The strains varied in their ability to cause lesions at the different concentrations, particularly the non-pathogen RMA1 which failed to induce more than limited browning (score 2), except when inoculated at the highest concentration. The *Pss* strain induced rapid lesion formation within 24 hours and on rare occassions spread slightly from the site of inoculation. At the higher concentrations the final lesions of all strains were similar in appearance, but could be differentiated by symptom timing. The timing of lesion onset was found to clearly differentiated the pathogenic *Psm* races from other strains. The non-pathogens (including the plum *Psm* R1-5300) and pathogenic strain *Pss* 9097 all induced rapid lesion formation at the highest concentration, with complete browning of the inoculation site (score 4) within the first 48 hours, which was suggestive of a HR. Pathogenic *Psm* R1 and R2 induced slower symptom development. This was indicative of a hemi-biotrophic interaction with the host. To study this interaction in more detail *Psm* R2 was inoculated onto detached leaves and electron microscopy used to examine bacteria-plant cell interactions. The bacteria were found to multiply initially in the apoplastic space without causing plant cell death, confirming hemi-biotrophic development, although some wall alterations were noted next to colonies. (Figure S9A -C).

To compare the host reactions on cherry and plum, the leaf population count and symptom scoring experiments were extended with a group of representative strains (Figure 8). As before, population counts clearly differentiated pathogens and non-pathogens (Figure 8A). On both hosts, pathogens exceeded 10^7^ CFU/ml and produced necrotic lesions. In comparison, non-pathogens failed to induce symptoms and did not reach 10^7^ CFU/ml. In the symptom scoring experiment (Figure 8B) all strains gave symptoms in the leaves, however the timing of symptoms was used to differentiate pathogenicity and hypersensitivity. On cherry, both *Pss* pathogens and the non-pathogens *Ps* 9643, R1-5300 and RMA1 induced symptoms rapidly, R2-leaf, *Pph* and *Psav* were slightly slower and R1-5244 only induced symptoms 48-72 hpi. In plum, the two *Pss* strains and *Ps* 9643 rapidly induced symptoms. Other non-pathogens were slower and not significantly faster than R2-leaf. Symptom development of plum *Psm* R1 5300 was not significantly different from cherry R1-5244, both inducing symptoms 48 hpi, indicating that in plum the two pathogens behave similarly. Representative images of symptoms on cherry and plum leaves over time are presented in Figure S10.

**Figure 8.**
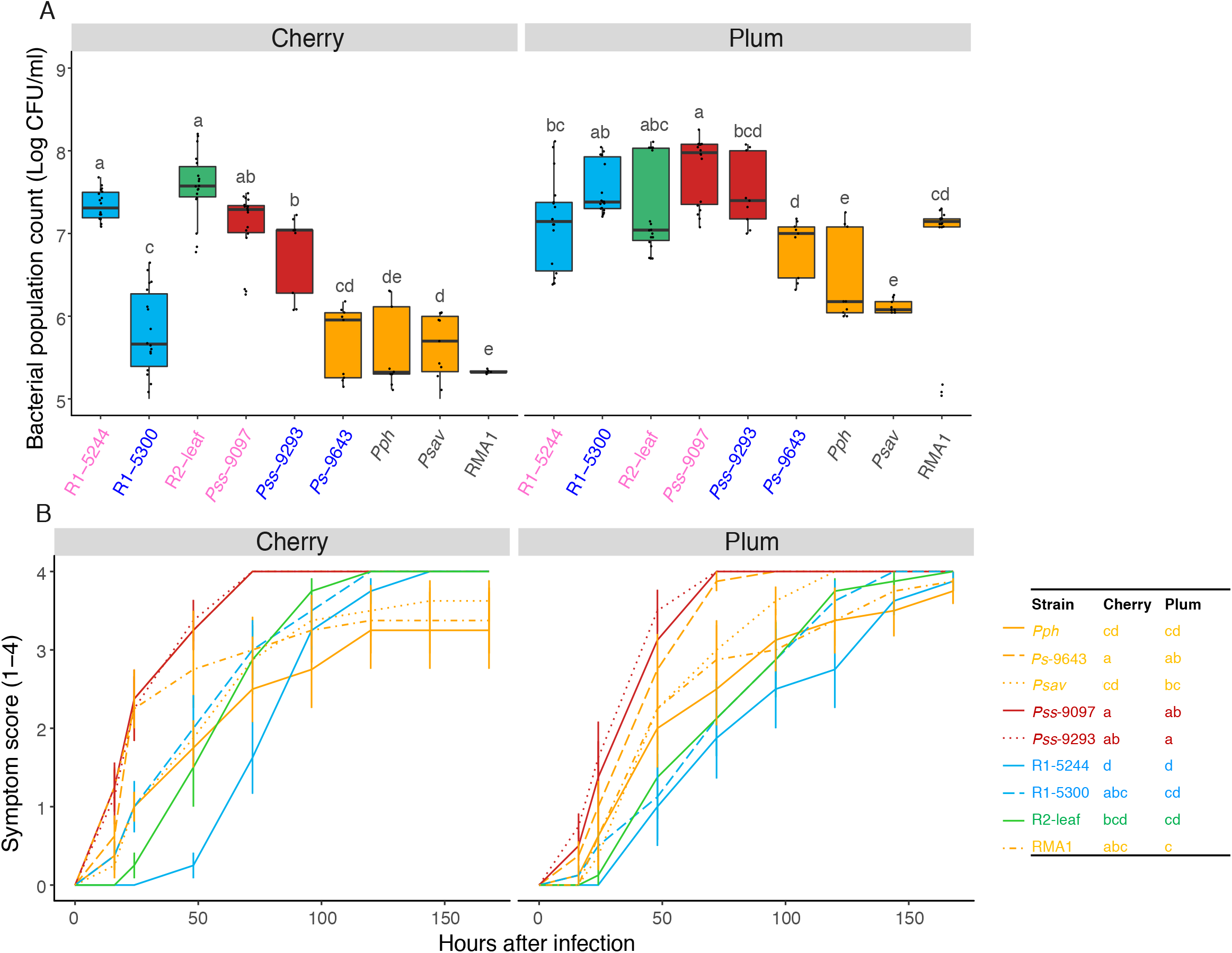
Pathogenicity of different strains, assessed by population counts and symptom scores, on cherry and plum leaves. A: Boxplots of day 10 population counts on cherry cv. Van and plum cv. Victoria. Strains isolated from cherry are pink whilst plum are blue. Boxplots are colour-coded by clade, with *Psm* R1: blue, *Psm* R2: green, *Pss:* red, non-pathogen: orange. Data presented are all the values for each treatment of two independent experiments (n=18). ANOVAs for both cherry and plum revealed significant differences between strains (p<0.01, df=8). Tukey-HSD (p=0.05, confidence level: 0.95) significance groups for the different strains are presented. B: Symptom development over time. Symptoms were scored, 0: no symptoms, 1: limited browning, 2: <50% inoculated area brown, 3: >50% inoculated area brown, 4: Complete browning. Strains are colour-coded as in A. Data presented are the mean values for each treatment of two independent experiments (n=8). Symptom development over time was analysed using the Area Under the Disease Progression Curve (AUDPC) analysis. ANOVAs for both cherry and plum revealed significant differences between strains (p<0.01, df=8). Tukey-HSD (p=0.05, confidence level: 0.95) significance groups are presented in the table next to the plot.

### Suitability of leaves for resistance screening

The leaf population assay clearly differentiated pathogens from non-pathogens. However, a screen for canker resistance would involve discriminating subtle differences in pathogen growth on different cherry genotypes. To see if detached leaves could discriminate cultivar differences, the assay was tested on four cultivars with differences in susceptibility recorded on woody tissue in the field. Strains representing the three cherry-infecting pathovars were tested. The three strains were able to grow to exceed 10^6^ CFU/ml (Figure 9) and cause disease symptoms in all cultivars. This suggested that on leaves, any host-resistance to the pathogens could not be easily discriminated. The leaf system, although useful for comparing strains with divergent virulence levels may not be sensitive enough to detect the subtle differences between races of the pathogens found in the field.

**Figure 9.**
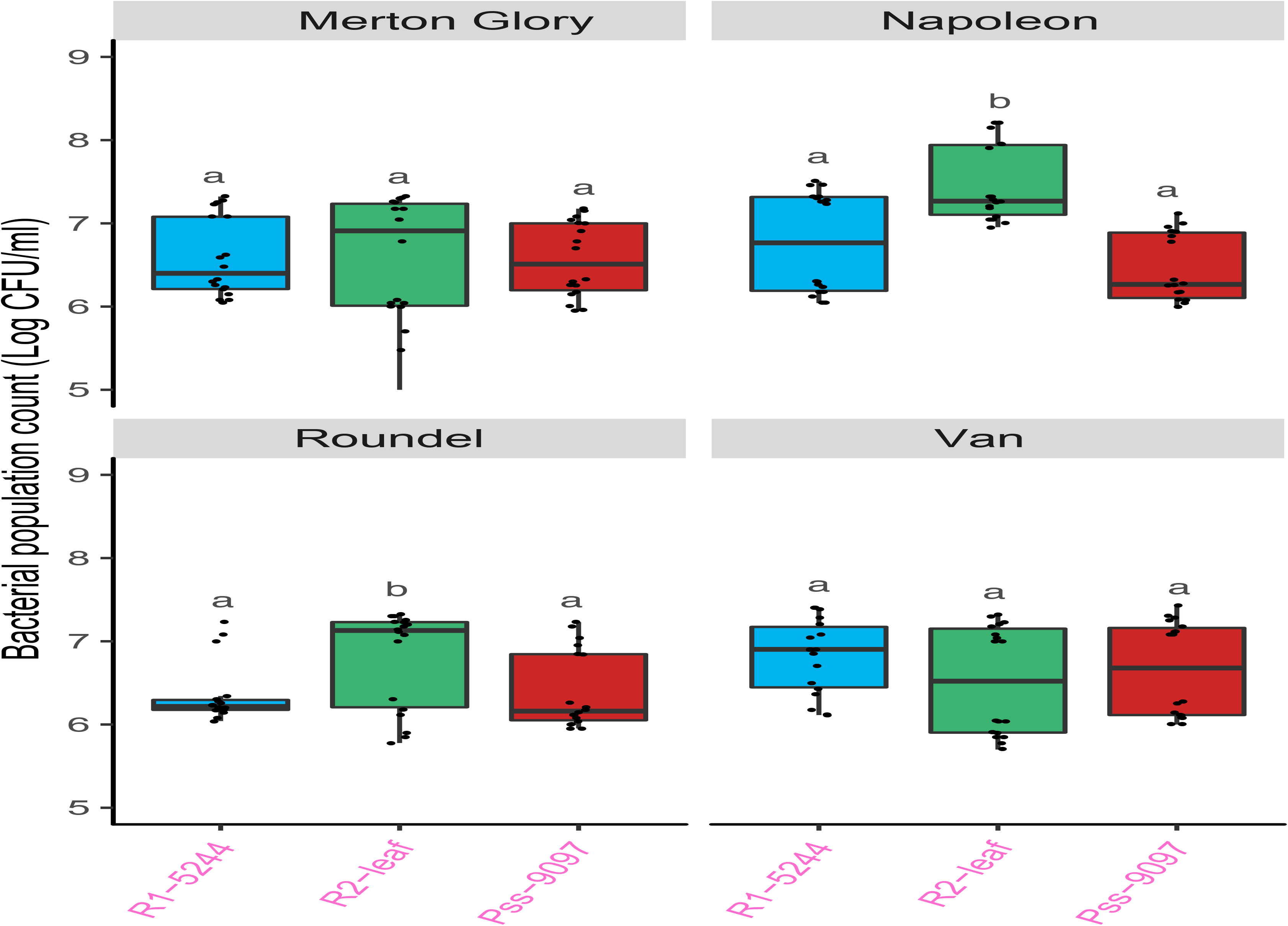
Bacterial multiplication recorded in different cherry cultivars. Boxplot of day 10 population counts of three pathogenic *P. syringae* strains on different cherry cultivars. Strains are colour-coded by clade, with *Psm* R1: blue, *Psm* R2: green, *Pss:* red. All three strains were cherry isolates so the names are coloured pink. Data presented are all the values for each treatment of two independent experiments (n=18). Tukey-HSD (p=0.05, confidence level: 0.95) significance groups for the different strains on each separate cultivar are presented. An ANOVA revealed significant differences between strains (p<0.01, df=2), cultivars (p<0.01, df=3) and a significant interaction (p<0.01, df=6).Tukey-HSD groups comparing the different cultivars are also presented.

## Discussion

In this study, the genomes of a set of *P. syringae* isolates from different hosts were sequenced. The ability of these strains to cause canker disease on cherry and plum was characterised. Breeding for resistance towards this complex disease is particularly challenging due to the large number of divergent strains that appear to be pathogenic. Host resistance to cherry canker is likely to be multi-factorial and potential mechanisms of resistance towards the different cherry-infecting clades may operate at different stages of the disease cycle. Phylogenetic analysis confirmed that the three major canker-causing clades *(Psm* R1, *Psm* R2 and *Pss)* fall in separate phylogroups, and therefore pathogenicity towards cherry has arisen multiple times in the *P. syringae* species complex. As the different clades have convergently evolved, it is likely that host resistance mechanisms targeted towards them differ significantly.

First, the ability of individual bacterial strains to cause cherry canker was assessed using a glasshouse whole-tree inoculation. This provided a baseline to compare with the results of laboratory-based assays, that may or may not correlate with ability to cause canker. Although strains of *Psm* R1 were phylogenetically indistinct, they could be divided into pathogenic and non-pathogenic isolates, with non-pathogenic isolates failing to cause gumming and black necrosis. Non-pathogenic strains isolated from distantly related plant species were unable to cause disease, supporting the theory that individual pathovars are mostly specialised to their particular host plant (Sarkar *et al*., 2006). All *Psm* R1 isolates from plum were non-pathogenic on cherry. Their lack of cherry pathogenicity may be due to host-specific factors. By contrast, all isolates of *Pss* (from plum and cherry) caused disease on cherry, indicating that these strains exhibit a greater host range.

Strains with variable virulence levels were then pathogenicity tested under field conditions, in assays which should be representative of natural disease. The different host-specificities of *Psm* R1 strains on cherry and plum were confirmed. The cherry isolate *Psm* R1 5244 was pathogenic to both cherry and plum, whereas *Psm* R1 5300 was only pathogenic on plum trees. This is an interesting result as phylogenetics revealed this clade to be highly homogeneous (Figure 1). As the phylogenetic analysis was based only on core house-keeping genes in the core genome, it may be missing divergence in the flexible genome that are responsible for differences in pathogenicity. Genomic analysis of these strains could reveal important differences in virulence factor repertoires that dictate host specificity. Interestingly, the results support studies done at East Malling looking at *Psm* R1 host specificity (Crosse & Garrett 1970). *Psm* R1 was originally designated as a race based on differences with *Psm* R2, however it is now known that these are two divergent clades, so should not really be designated as races of the same pathovar. However, the differences in pathogenicity of members of *Psm* R1 may indicate that, at least within the bacterial populations occupying orchards in UK, there may be a race structure within this clade, with the different groups varying in ability to infect different *Prunus* species. Members of the group containing *Psm* R1-5300 may be restricted in growth on cherry due to the expression of avirulence factors. Further sampling of a diverse range of strains from different *Prunus* species and cultivars should confirm this hypothesis.

The field inoculations were assessed using both disease score and symptom length. For disease score, cherry leaf scars appeared much less susceptible to infection. The leaf scar may act as a barrier to infection and reduce bacterial concentrations as the bacterial population is bottle-necked. Therefore, a higher percentage of trees scored highly for wound inoculations as this by-passed the barrier to infection. For symptom lengths, results were more variable, with most pathogen inoculations only spreading slightly. Only in rare cases did they cause severe necrosis, sometimes exceeding 100mm. The results revealed significant differences between cherry cultivars. In both wound and leaf scar inoculations, the cultivar Merton Glory exhibited a broad level of tolerance to all three pathogenic clades. This cultivar is therefore a candidate for further study of the mechanisms underlying resistance. Although the analysis did not show a strain by cultivar interaction, there was variation in resistance to *Psm* R2. This strain was associated with only limited disease on Van, whilst the cultivar Roundel was highly susceptible. Van is therefore a candidate cultivar exhibiting race-specific resistance. There was no significant difference in symptom length or disease score of pathogenic *Psm* R1 and R2 on Napoleon, contrasting to previous studies that suggested Napoleon to be resistant to R2 (Garrett, 1978). In addition, previous studies reported *Pss* and *Psm* R2 to be less invasive through leaf scars inoculations than *Psm* R1 (Crosse & Garrett 1966; Freigoun & Crosse 1975), which contrasts to this study where all clades caused disease. Differences in experimental procedure could have led to variation in results. The original studies used fully mature trees which may exhibit contrasting resistance mechanisms to the young trees used in this study (Freigoun & Crosse 1975; Garrett 1978).

The field experiment on plum demonstrated significant differences between strains, but not between cultivars. The cultivar Marjorie’s Seedling was not susceptible to any strains inoculated through leaf scars indicating this is unlikely to be a natural entry point for pathogens. Indeed, previous reports suggest that plum pathogens do not naturally enter through the leaf scars (Crosse, 1966). However, some pathogenic strains were found to be capable of causing disease on the cultivar Victoria when inoculated through leaf scars.

Several rapid laboratory-based assays were tested for their suitability for resistance screening. Assuming that the field wound inoculations represent the natural disease, the results of the other tests for a set of strains on cherry cv. Van were correlated against the wound results (Figure 10). The leaf scar, cut shoot and leaf population assays all correlated well with the wound results (r > 0.70), whilst the fruit assay did not correlate well (r = 0.37).

**Figure 10.**
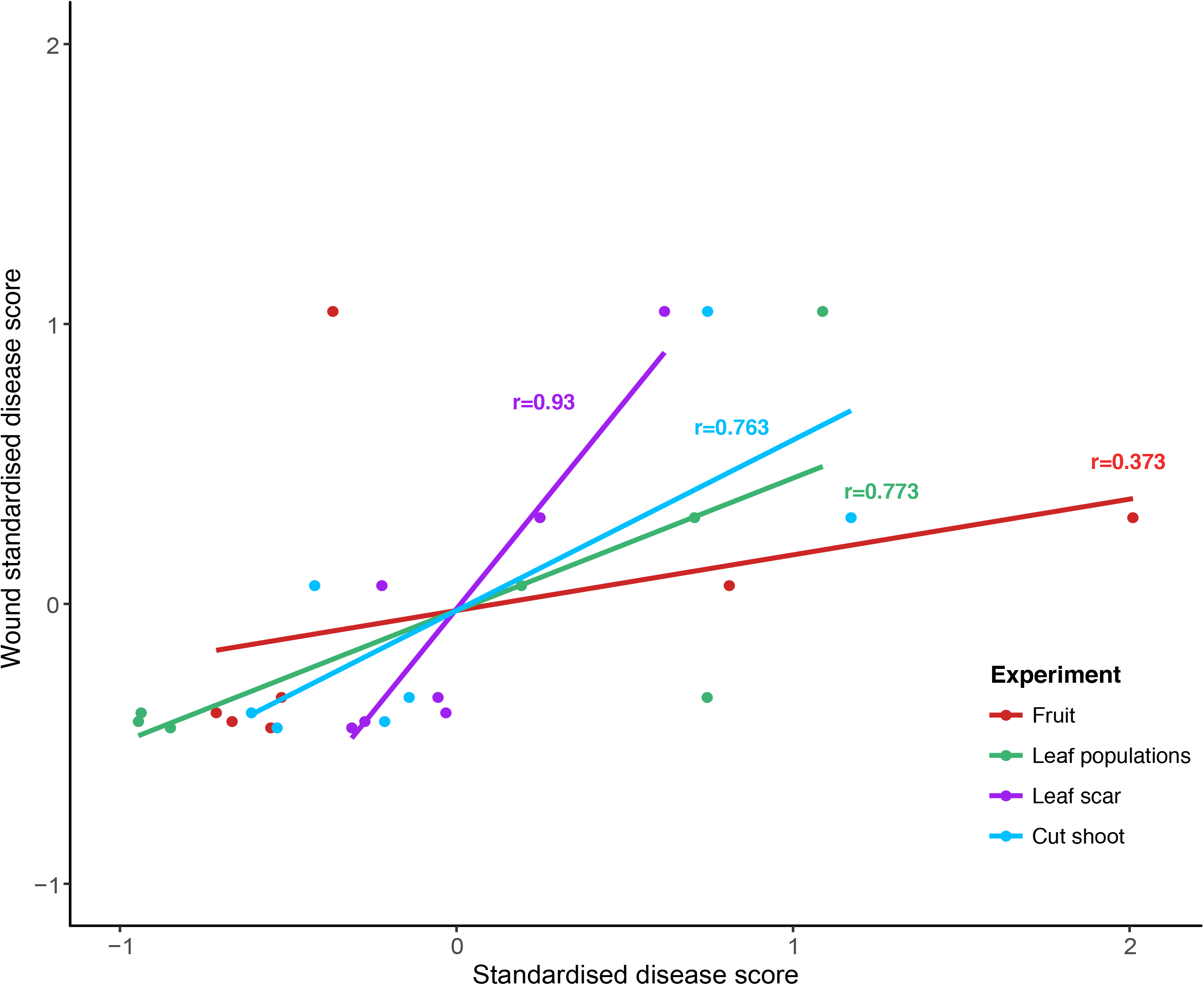
Correlation of different inoculation experiments with the whole-tree wound inoculations performed in the field. The scatterplot was created using the mean standardised disease scores for each bacterial strain on cherry cv. Van. A linear model (lm) line was plotted for each experiment. The Pearson’s correlation coefficients correlating the results of each experiment with the wound field inoculations are presented.

Both inoculations of woody tissues (leaf scar and cut shoots) correlated well with the wound results. The cut shoot assay provided a rapid assessment that could differentiate pathogens and nonpathogens. It was also sensitive enough to detect variation in pathogen virulence on the different cultivars. For example, *Psm* R2 caused the greatest necrosis on cv. Roundel in all three woody tissue inoculation tests, indicating that this cultivar is highly susceptible to *Psm* R2. In addition, differential virulence of *Psm* R1 and *Psm* R2 on cv. Van was supported by both the cut shoot and field experiments, suggesting this cultivar may possess some resistance to *Psm* R2. Various studies have utilised cut shoot inoculations of *P. syringae* and fungal canker-causing pathogens. Differences in virulence of the same isolate between field and laboratory results are sometimes reported (Farhadfar *et al*., 2016; Gomez-Cortecero *et al*., 2016). Therefore, a combination of whole tree and shoot tests could provide the most robust method of assessment. The cut shoot assay provided a means to perform rapid high-throughput screening, with speed aided through automated image analysis of shoots (Li *et al*., 2015). Therefore, the results of such tests may help narrow down a list of putatively resistant genotypes, before resistance testing on whole trees.

The lack of correlation of the cherry fruit test with the field experiment indicated that the lesion development on fruit induced by a *P. syringae* strain may not reflect its pathogenicity in the field. Nevertheless, the qualitative symptoms this assay provides are useful to rapidly differentiate the different pathogenic clades. The induction of symptoms by strains non-pathogenic in the field (e.g. *Ps* 9643, Figure S4), indicated that results must be considered with caution. In comparison, the leaf population assay correlated well with wound inoculations and allowed discrimination of pathogens and non-pathogens. However, when the three pathogenic clades were inoculated across cherry cultivars (Figure 9) they all exceeded 10^6^ CFU/ml *in planta* and caused symptom development. The presence of similar symptoms on both susceptible and tolerant varieties means that this method would not be very applicable for large-scale screening. The leaf and fruit assays were therefore not sensitive enough to determine subtle differences in cultivar susceptibility seen in the field experiment such as the resistance of cherry cv. Van to Psm R2. The quantitative differences in resistance of the different cherry cultivars in the field and shoot experiments may be tissue-specific, and therefore resistance phenotypes in fruit and leaves may differ substantially from those found in dormant woody tissues.

Detached leaves provided a rapid means to assess pathogenicity through the measurement of bacterial population counts over time. Cherry leaf population counts clearly discriminated pathogenic and non-pathogenic strains. When inoculated at a low concentration only pathogenic strains (R1-5244, R1-leaf and *Pss* 9097) were able to cause disease lesions on cherry, which appeared 7-10 dpi. On Plum, the non-pathogen RMA1 and plum isolate R1-5300 were able to grow to similar levels to the cherry pathogens (Figure 7). The fact that RMA1 was able to grow to high levels in plum leaves does not correspond to its pathogenicity in the field assay. Interestingly, in the cutshoot assay (Figure 5) RMA1 caused necrosis on plum similar to the *Pss* pathogen. The field experiment showed that RMA1 is not a true pathogen of plum, however, its virulence in the lab-based assays may indicate it has adaptive potential to cause disease when inoculated in unnaturally high concentrations directly onto plant tissue. Its inability to cause any disease on cherry in all lab-based assays indicated that cherry may exhibit a robust non-host immune response towards this non-pathogen, which is different to that expressed in plum.

Symptom development on cherry leaves allowed differentiation of hypersensitive and pathogenic responses. When inoculated at high concentrations all strains produced necrotic lesions, however non-pathogens were found to induce symptoms earlier than pathogens of *Psm* R1 and R2. The activation of the HR may mean that ETI is operating against non-pathogens in cherry leaves, and differences in effector repertoires between cherry-infecting strains and non-pathogens could reveal those effectors that are detected. In particular, there were clear differences in pathogenicity of the two *Psm* R1 strains on cherry, which agreed with the whole-tree assay. The HR on cherry was clear for non-pathogens *Psm* R1 5300, *Ps* 9643 and RMA1, whereas symptom development associated with *Pph* and *Psav* was slower. This slower onset of symptoms may mean that any hypersensitive response induced by these strains is weaker or that more basal resistance mechanisms such as PAMP-triggered immune responses play a greater role in preventing their population growth in leaves. Interestingly, although *Pss* strains reached high population levels in the leaves, they triggered symptom development at a similar rate to the HR caused by non-pathogens. *P. syringae* is traditionally described as a hemi-biotrophic pathogen (Lindeberg *et al*., 2012), with delayed symptom onset during the biotrophic phase followed by symptoms during a necrotrophic phase. The results indicated that on leaves *Pss* may be more necrotrophic as it triggers symptoms rapidly. Further study could reveal the factors inducing these rapid symptoms. The production of non-ribosomal peptide toxins is common in strains of phylogroup 2, which includes *Pss* (Dudnik & Dudler, 2014), and if expressed early could cause the necrotic symptoms seen. Indeed, a study of *Pss* toxins (Yin-Yuan & Gross, 1991) showed that syringomycin is expressed within the first 24 hours of inoculation of immature cherry fruits. *Pss* could also be deliberately triggering the HR like other necrotrophic pathogens to aid disease development (Govrin & Levine, 2000). Further study of the immune responses occurring within plant cells would be required to test these hypotheses.

The failure of designated non-pathogenic strains to produce symptoms in woody tissues was reflected by their low multiplication and induction of a HR-like response in leaves. Such clear cut resistance is characteristic of *avr/R* gene interactions reflecting ETI. By contrast, where differential reactions were observed between cultivars challenged with pathogenic *Psm* and *Pss*, quantitative differences in symptoms were seen. The lack of clear differentials between cultivars suggests that variation in susceptibility is not based simply on *avr/R* gene recognition. In field conditions, this plant-pathogen interaction lasts for many months. Perhaps, factors important not just for pathogenicity, but for the ability of bacterial populations to successfully colonise and persist through the season, dictate the outcome of this interaction. Resistance mechanisms that reduce persistence in woody tissue, e.g. responses that block bacteria spreading to new tissues and acquiring nutrients, may prevent a pathogenic strain from causing severe disease.

This study has focused on the detailed analysis of pathogenicity in strains used for genome sequencing. Results show that representatives of the three clades of *P. syringae* that cause bacterial canker may utilise distinct mechanisms of virulence and trigger differing host resistance mechanisms in cherry. A HR is putatively triggered in leaves, indicating that effector-triggered immunity may be operating in cherry against pathogens of other hosts. Cherry leaves and fruit failed to sufficiently reveal varietal differences to the same extent as experiments on woody tissues. This suggests that some resistance mechanisms are tissue-specific. A whole range of complex variable traits could be involved in these varietal differences in susceptibility. These include timing of leaf drop, phellogen activity and differences in leaf-surface bacterial populations which act as inocula for wood infections, as discussed by Crosse (1966). Breeding resistance to at least three rather distinct groups of a pathogen remains a challenging prospect. Cultivars such as Merton Glory that exhibit resistance to all three clades may be useful for determining the genetic basis of broad-spectrum resistance mechanisms, independent of ETI.

## Acknowledgments

We thank Steve Roberts, Helen Neale and David Guttman for providing bacterial strains used in this study. Electron microscopy was performed by Dr Ian Brown (University of Kent). We also thank Karen Russell and Connie Garrett for valuable advice about the pathogenicity test development, as well as the East Malling Farm and Glass team for assistance in tree grafting and planting. The authors declare no conflict of interest.

## Supporting material

**Figure S1**

Images of immature cherry fruits inoculated with *Psm* R1 strains. Images were taken 10dpi. Five replicate cherries were inoculated per strain. Strains are colour-coded based on host of isolation as pink (cherry) or blue (plum).

**Figure S2**

Images of immature cherry fruits inoculated with *Psm* R2 strains. Images were taken 10dpi. Five replicate cherries were inoculated per strain. Strains are colour-coded based on host of isolation as pink (cherry).

**Figure S3**

Images of immature cherry fruits inoculated with *Pss* strains. Images were taken 10dpi. Five replicate cherries were inoculated per strain. Strains are colour-coded based on host of isolation as pink (cherry) or blue (plum).

**Figure S4**

Images of immature cherry fruits inoculated with non-pathogen strains and a no-bacteria control. Images were taken 10dpi. Five replicate cherries were inoculated per strain.

**Figure S5**

Boxplot of diameter of necrosis caused by cherry pathogens on four cherry cultivars using immature green cherry fruits. Strains are colour-coded with those isolated from cherry in pink and the no bacterial control in black. The boxplots are colour-coded by clade: *Psm* R1: blue, *Psm* R2: green, *Pss:* red and no bacteria control: black. Data presented are all values (n=20) per treatment of two independent experiments. An ANOVA revealed significant differences between strains (p<0.01, df=3), cultivars (p<0.01, df=3) and a significant interaction (p<0.01, df=9). Tukey-HSD (p=0.05, confidence level: 0.95) significance groups for the different strains for each separate cultivar are presented above each boxplot.

**Figure S6**

Symptoms observed in detached cherry leaves using different inoculation methods. Representative images of the four methods – infiltration, stab, droplet and wound + droplet. Leaves show inoculation with *Psm* R1-5244 or a 10mM MgCl2 control.

**Figure S7**

Boxplot of day 10 population counts of all strains used in this study on cherry cv. Van leaves. Strains are colour-coded with those isolated from cherry in pink, plum in blue and non-pathogens in black. The boxplots are coloured by clade: *Psm* R1: blue, *Psm* R2: green, *Pss:* red. The 10mM MgCl2 control is not included as no bacteria were found. The data presented are all values for each treatment (n=9). An ANOVA revealed significant differences between strains (p<0.01, df=20). Tukey-HSD (p=0.05, confidence level: 0.95) significance groups for the different strains are presented.

**Figure S8**

Symptom development over time after inoculation of various *P. syringae* strains in cherry cv. Van at different concentrations. Strains are colour-coded, with those isolated from cherry in pink and plum in blue. Symptoms were scored from 0-5. 0: no symptoms, 1: limited browning, 2: browning <50% of inoculated site, 3: browning >50% of inoculated site, 4: Complete browning, 5: Spread from site of inoculation. Data presented are the means (n=4) and error bars show the standard error above and below the mean. The lines for each strains are colour-coded with *Psm* R1: blue, *Psm* R2: green, *Pss:* red, non-pathogen RMA1: orange. Symptom development over time was analysed using AUDPC. An ANOVA revealed significant differences between strains (p<0.01, df=4), concentrations (p<0.01, df=3) and a significant interaction (p<0.01, df=12). Tukey-HSD (p=0.05, confidence level: 0.95) significance groups are presented.

**Figure S9**

TEM images of *Psm* R2-leaf in a detached cherry leaf one week after inoculation. Arrows point to putative papilla formation in the plant cell wall next to a bacterial colony containing dead bacterial cells. A: Bacteria inhabiting apoplastic space next to cells. Note that no cell wall alterations appeared in the plant cells. B: Cell wall alterations (papilla formation) shown by arrows in plant cells. C: A bacterial colony containing dead and alive bacteria next to plant cells.

**Figure S10**

Images of symptom development over time on cherry and plum. A: Cherry cv. Van, B: Plum cv. Victoria. The same leaf was imaged 16, 24, 48 and 72hpi. Arrows indicate the first appearance of symptoms for that particular strain. Strains are labelled: 1: *Psm* R1-5244, 2: *Psm* R1-5300, 3: *Psm* R2-leaf, 4: Ps-9643, 5: Pss-9097, 6: Pss-9293, 7: RMA1, 8: *Psav*, 9: *Pph*, C: No bacteria control

**Table S1**

Proportional Odds Model (POM) analysis of the glasshouse whole-tree wound inoculations. Model comparisons are first shown with the ANOVA comparing models. The summary of the final model (score ~g1) is shown along with lsmeans Tukey-HSD groupings of strains (corresponds to groupings on Figure 2).

**Table S2**

REML analysis of field inoculation of cherry inoculated by leaf scar. The REML model and ANOVA are presented, followed by lsmean Tukey-HSD groupings for cultivars, strains and then strains on each cultivar (corresponds to groupings on Figure 3).

**Table S3**

REML analysis of field inoculation of cherry inoculated by wound. The REML model and ANOVA are presented, followed by lsmean Tukey-HSD groupings for cultivars, strains and then strains on each cultivar (corresponds to groupings on Figure 3).

**Table S4**

REML analysis of field inoculation of plum inoculated by leaf scar. The REML model and ANOVA are presented, followed by lsmean Tukey-HSD groupings for strains and then strains on each cultivar (corresponds to groupings on Figure 4).

**Table S5**

REML analysis of field inoculation of plum inoculated by wound. The REML model and ANOVA are presented, followed by lsmean Tukey-HSD groupings for strains and then strains on each cultivar (corresponds to groupings on Figure 4).

**Table S6**

POM analysis of the cherry field inoculations. Model comparisons are first shown with the ANOVA comparing models. The summary of the final model (score~strain+cv+ino+block) is then presented.

**Table S7**

Lsmean Tukey-HSD groupings for different treatment combinations from the POM analysis of cherry field inoculations. Groups for strains on different cultivars are presented (corresponds to groupings on Figure 3), followed by groupings of cultivars in each inoculation method and then strains across the two inoculation methods.

**Table S8**

POM analysis of the plum field inoculations. Model comparisons are first shown with the ANOVA comparing models. The summary of the final model (score~strain+cv+ino+block) is then presented.

**Table S9**

Lsmeans Tukey-HSD groupings for different treatment combinations from the POM analysis of plum field inoculations. Groups for strains on different cultivars are presented (corresponds to groupings on Figure 3), followed by groupings of cultivars in each inoculation method and then strains across the two inoculation methods.

**Table S10**

ANOVA table of cut shoot inoculations followed by lsmeans Tukey-HSD groupings for the strains on each cultivar (corresponds to groupings on Figure 5).

**Table S11**

ANOVA table of immature cherry fruit inoculations of all isolates followed by Tukey-HSD groupings for the strains extracted using the agricolae package function HSD.test (corresponds to groupings on Figure 6).

**Table S12**

REML analysis of immature cherry fruit inoculations where different bacterial strains were inoculated onto different host cultivars. The REML model is presented. Lsmeans Tukey-HSD groups for strains on different cultivars are presented (corresponds to groupings on Figure S5), followed by groupings based on all possible treatments.

**Table S13**

ANOVA table of day 10 leaf population counts of different bacterial strains inoculated on cherry and plum. Tukey-HSD groups for strains are presented (corresponds to groupings on Figure 7).

**Table S14**

REML analysis of day 10 leaf population counts of reference bacterial strains inoculated on cherry leaves. The model is shown followed by ANOVA table. Lsmeans Tukey-HSD groups for strains are presented (corresponds to groupings on Figure 8).

**Table S15**

REML analysis of day 10 leaf population counts of reference bacterial strains inoculated on plum leaves. The model is shown followed by ANOVA table. Lsmeans Tukey-HSD groups for strains are presented (corresponds to groupings on Figure 8).

**Table S16**

ANOVA table of AUDPC analysis of leaf symptom score over time of different bacterial strains inoculated on cherry. Tukey-HSD groups for strains are presented (corresponds to groupings on Figure 8).

**Table S17**

ANOVA table of AUDPC analysis of leaf symptom score over time of different bacterial strains inoculated on cherry. Tukey-HSD groups for strains are presented (corresponds to groupings on Figure 8).

**Table S18**

ANOVA table of day 10 leaf population counts of different bacterial strains inoculated on different cherry cultivars. Tukey-HSD groups for strains are presented (corresponds to groupings on Figure 9).

**Table S19**

ANOVA table of leaf population counts of all isolates used in this study, followed by Tukey-HSD groupings for the strains extracted using the agricolae package function HSD.test (corresponds to groupings on Figure S7).

**Table S20**

ANOVA table of AUDPC analysis of symptom score on leaves of several bacterial strains inoculated at different concentrations. This is followed by lsmeans Tukey-HSD groupings for the strains (corresponds to groupings on Figure S8).

